# Generalization guides human exploration in vast decision spaces

**DOI:** 10.1101/171371

**Authors:** Charley M. Wu, Eric Schulz, Maarten Speekenbrink, Jonathan D. Nelson, Bjöorn Meder

## Abstract

From foraging for food to learning complex games, many aspects of human behaviour can be framed as a search problem with a vast space of possible actions. Under finite search horizons, optimal solutions are generally unobtainable. Yet how do humans navigate vast problem spaces, which require intelligent exploration of unobserved actions? Using a variety of bandit tasks with up to 121 arms, we study how humans search for rewards under limited search horizons, where the spatial correlation of rewards (in both generated and natural environments) provides traction for generalization. Across a variety of different probabilistic and heuristic models, we find evidence that Gaussian Process function learning—combined with an optimistic Upper Confidence Bound sampling strategy—provides a robust account of how people use generalization to guide search. Our modelling results and parameter estimates are recoverable, and can be used to simulate human-like performance, providing insights about human behaviour in complex environments.

## Introduction

Many aspects of human behaviour can be understood as a type of search problem^1^, from foraging for food or resources^2^, to searching through a hypothesis space to learn causal relationships^3^, or more generally, learning which actions lead to rewarding outcomes^4^. In a natural setting, these tasks come with a vast space of possible actions, each corresponding to some reward that can only be observed through experience. In such problems, one must learn to balance the dual goals of exploring unknown options, while also exploiting familiar options for immediate returns. This frames the *exploration-exploitation dilemma*, typically studied using the multi-armed bandit problems^5–8^, which imagine a gambler in front of a row of slot machines, learning the reward distributions of each option independently. Solutions to the problem propose different policies for how to learn about which arms are better to play (exploration), while also playing known high-value arms to maximize reward (exploitation). Yet under real-world constraints of limited time or resources, it is not enough to know *when* to explore; one must also know *where* to explore.

Human learners are incredibly fast at adapting to unfamiliar environments, where the same situation is rarely encountered twice^9, 10^. This highlights an intriguing gap between human and machine learning, where traditional approaches to reinforcement learning typically learn about the distribution of rewards for each state independently^4^. Such an approach falls short in more realistic scenarios where the size of the problem space is far larger than the search horizon, and it becomes infeasible to observe all possible options^11, 12^. What strategies are available for an intelligent agent—biological or machine—to guide efficient exploration when not all options can be explored?

One method for dealing with vast state spaces is to use *function learning* as a mechanism for generalizing prior experience to unobserved states^13^. The function learning approach approximates a global value function over all options, including ones not experienced yet^10^. This allows for generalization to vast and potentially infinite state spaces, based on a small number of observations. Additionally, function learning scales to problems with complex sequential dynamics and has been used in tandem with restricted search methods, such as Monte Carlo sampling, for navigating intractably large search trees^14, 15^. While restricted search methods have been proposed as models of human reinforcement learning in planning tasks^16, 17^, here we focus on situations in which a rich model of environmental structure supports learning and generalization^18^.

Function learning has been successfully utilized for adaptive generalization in various machine learning applications^19, 20^, although relatively little is known about how humans generalize *in vivo* (e.g., in a search task; but see^8^). Building on previous work exploring inductive biases in pure function learning contexts^21, 22^ and human behaviour in univariate function optimization^23^, we present a comprehensive approach using a robust computational modelling framework to understand how humans generalize in an active search task.

Across three studies using uni-and bivariate multi-armed bandits with up to 121 arms, we compare a diverse set of computational models in their ability to predict individual human behaviour. In all experiments, the majority of subjects are best captured by a model combining function learning using Gaussian Process (GP) regression, with an optimistic Upper Confidence Bound (UCB) sampling strategy that directly balances expectations of reward with the reduction of uncertainty. Importantly, we recover meaningful and robust estimates about the nature of human generalization, showing the limits of traditional models of associative learning^24^ in tasks where the environmental structure supports learning and inference.

The main contributions of this paper are threefold:

1. We introduce the *spatially correlated multi-armed bandit* as a paradigm for studying how people use generalization to guide search in larger problems space than traditionally used for studying human behaviour.
2. We find that a Gaussian Process model of function learning robustly captures how humans generalize and learn about the structure of the environment, where an observed tendency towards undergeneralization is shown to sometimes be beneficial.
3. We show that participants solve the exploration-exploitation dilemma by optimistically inflating expectations of reward by the underlying uncertainty, with recoverable evidence for the separate phenomena of directed (towards reducing uncertainty) and undirected (noisy) exploration.

## Results

A useful inductive bias in many real world search tasks is to assume a spatial correlation between rewards^25^ (i.e., clumpiness of resource distributions^26^). This is equivalent to assuming that similar actions or states will yield similar outcomes. We present human data and modelling results from three experiments (Fig. 1) using univariate (Experiment 1) and bivariate (Experiment 2) environments with fixed levels of spatial correlations, and also real-world environments where spatial correlations occur naturally (Experiment 3). The spatial correlation of rewards provides a context to each arm of the bandit, which can be learned and used to generalize to not-yet-observed options, thereby guiding search decisions. Additionally, since recent work has connected both spatial and conceptual representations to a common neural substrate^27^, our results in a spatial domain provide potential pathways to other search domains, such as contextual^28–30^ or semantic search^31, 32^.

**Figure 1.**
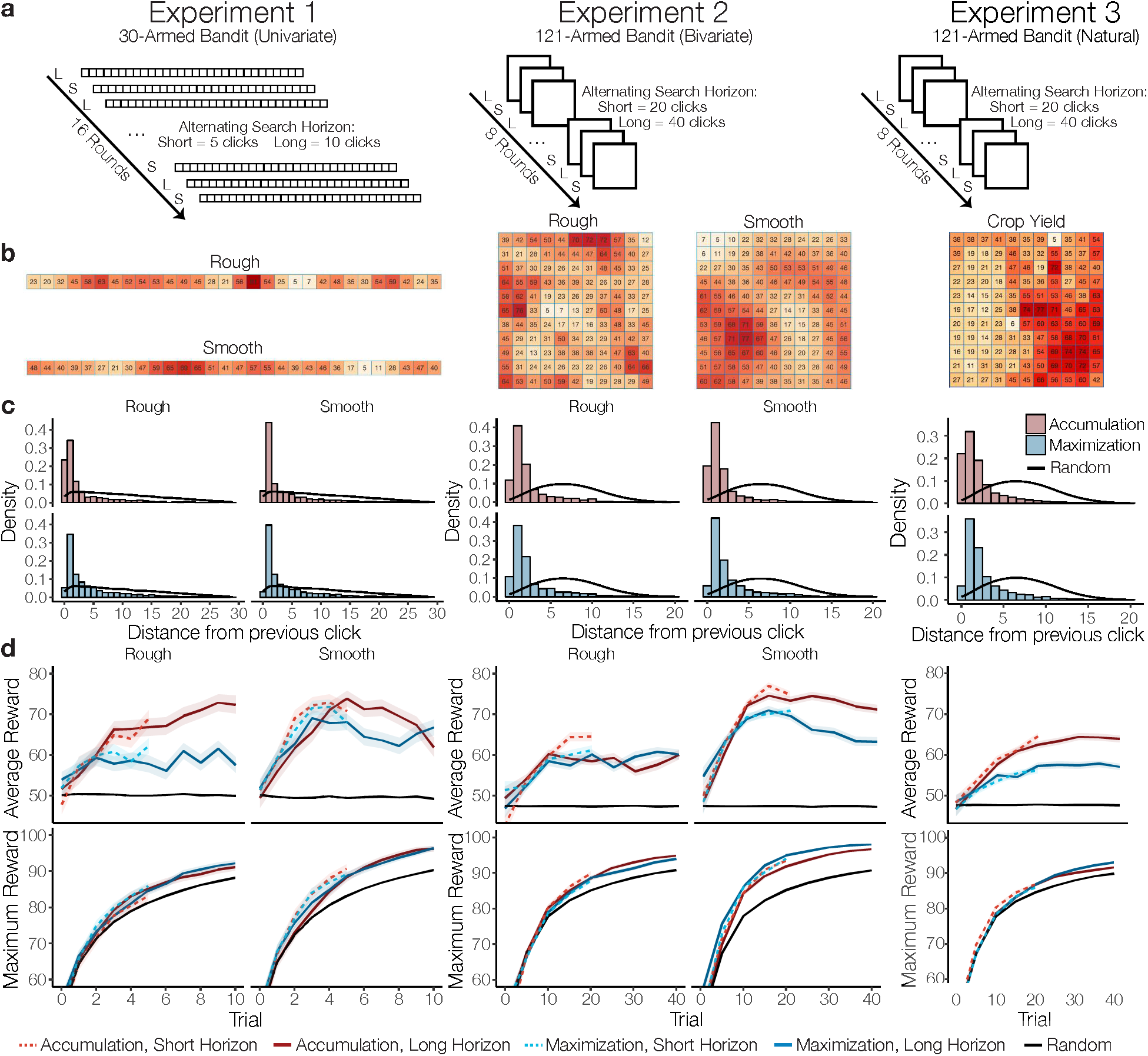
Procedure and behavioural results. Experiments 1 and 2 used a 2×2 between-subject design, manipulating the type of environment (Rough or Smooth) and the payoff condition (Accumulation or Maximization), while Experiment 3 manipulated only payoff conditions (between subjects) and used a set of natural environments where rewards reflect normalized crop yields from various agricultural datasets. **a)** Experiment 1 used a 1D array of 30 possible options, while Experiments 2 and 3 used a 2D array (11×11) with 121 options. Experiments took place over 16 (Exp. 1) or 8 rounds (Exp. 2 and 3), with a new environment sampled without replacement for each round. Search horizons alternated between rounds (within subject), with the horizon order counter-balanced between subjects. **b)** Examples of fully revealed search environments, where tiles were initially blank at the beginning of each round, except for a single randomly revealed tile. Rough and Smooth environments differed in the extent of spatial correlations, while Crop Yield environments have no fixed level of correlation (see SI).**c)** Locality of sampling behaviour compared to a random baseline simulated over 10,000 rounds (black line), where distance is measured using Manhattan distance and the y-axis indicates the probability density of different distances (with a different maximum range for Exp. 1 compared to Exp. 2 and 3). **d)** Average reward earned (Accumulation goal) and maximum reward revealed (Maximization goal), where coloured lines indicate the assigned payoff condition and shaded regions show the standard error of the mean. Short horizon trials are indicated by lighter colours and dashed lines, while black lines are a random baseline simulated over 10,000 rounds.

### Experiment 1

Participants (*n* = 81) searched for rewards on a 1 × 30 grid world, where each tile represented a reward-generating arm of the bandit (Fig. 1a). The mean rewards of each tile were spatially correlated, with stronger correlations in *Smooth* than in *Rough* environments (between subjects; Fig. 1b). Participants were either assigned the goal of accumulating the largest average reward (*Accumulation condition*), thereby balancing exploration-exploitation, or of finding the best overall tile (*Maximization condition*), an exploration goal directed towards finding the global maximum. Additionally, the search horizons alternated between rounds (within subject; *Short* = 5 vs. *Long* = 10), with the order counter-balanced between subjects. We hypothesized that if function learning guides search behaviour, participants would perform better and learn faster in smooth environments, in which stronger spatial correlations reveal more information about nearby tiles^33^.

Looking first at sampling behaviour, the distance between sequential choices was more localized than chance (*t*(80) = 39.8, *p* < .001, *d* = 4.4, 95% CI (3.7,5.1), *BF* > 100; Fig. 1c; all reported *t*-tests are two-sided), as has also been observed in semantic search^31^ and causal learning^3^ domains. Participants in the Accumulation condition sampled more locally than those in the Maximization condition (*t*(79) = 3.33, *p* = .001, *d* = 0.75, 95% CI (0.3,1.2), *BF* = 24), corresponding to the increased demand to exploit known or near-known rewards. Comparing performance in different environments, the learning curves in Fig. 1d show that participants in Smooth environments obtained higher average rewards than participants in Rough environments (*t*(79) = 3.58, *p* < .001, *d* = 0.8, 95% CI (0.3,1.3), *BF* = 47.4), consistent with the hypothesis that spatial patterns in the environment can be learned and used to guide search. Surprisingly, longer search horizons (solid vs. dashed lines) did not lead to higher average reward (*t*(80) = 0.60, *p* = .549, *d* = 0.07, 95% CI (–0.4,0.5), *BF* = 0.2). We analyzed both average reward and the maximum reward obtained for each subject, irrespective of their payoff condition (Maximization or Accumulation). Remarkably, participants in the Accumulation condition performed best according to both performance measures, achieving higher average rewards than those in the Maximization condition (*t*(79) = 2.89, *p* = .005, *d* = 0.7, 95% CI (0.2,1.1), *BF* = 7.9), and performing equally well in terms of finding the largest overall reward (*t*(79) = –0.73, *p* = .467, *d* = –0.2, 95% CI (–0.3,0.6), *BF* = 0.3). Thus, a strategy balancing exploration and exploitation—at least for human learners—may achieve the global optimization goal *en passant*.

### Experiment 2

Experiment 2 had the same design as Experiment 1, but used a 11 × 11 grid representing an underlying bivariate reward function (Fig. 1 centre) and longer search horizons to match the larger search space (*Short* = 20 vs. *Long* = 40). We replicated the main results of Experiment 1, showing participants (*n* = 80) sampled more locally than a random baseline (*t*(79) = 50.1, *p* < .001, *d* = 5.6, 95% CI (4.7,6.5), *BF* > 100; Fig. 1c), Accumulation participants sampled more locally than Maximization participants (*t*(78) = 2.75, *p* = .007, *d* = 0.6, 95% CI (0.2,1.1), *BF* = 5.7), and participants obtained higher rewards in Smooth than in Rough environments (*t*(78) = 6.55, *p* < .001, *d* = 1.5, 95% CI (0.9,2.0), *BF* > 100; Fig. 1d). For both locality of sampling and the difference in average reward between environments, the effect size was larger in Experiment 2 than in Experiment 1. We also replicated the result that participants in the Accumulation condition were as good as participants in the Maximization condition at discovering the largest reward values (*t*(78) = –0.62, *p* = .534, *d* = –0.1, 95% CI (–0.6,0.3), *BF* = 0.3), yet in Experiment 2 the Accumulation condition did not lead to substantially better performance than the Maximization condition in terms of average reward (*t*(78) = –1.31, *p* = .192, *d* = –0.3, 95% CI (–0.7,0.2), *BF* = 0.5). Again, short search horizons led to the same level of performance as longer horizons, (*t*(79) = –0.96, *p* = .341, *d* = –0.1, 95% CI (–0.3,0.1), *BF* = 0.2), suggesting that learning occurs rapidly and peaks rather early.

### Experiment 3

Experiment 3 used the same 121-armed bivariate bandit as Experiment 2, but rather than generating environments with fixed levels of spatial correlations, we sampled environments from 20 different agricultural datasets^34^, where payoffs correspond to the normalized yield of various crops (e.g., wheat, corn, and barley). These datasets have naturally occurring spatial correlations and are naturally segmented into a grid based on the rows and columns of a field, thus requiring no interpolation or other transformation except for the normalization of payoffs (see SI for selection criteria). The crucial difference compared to Experiment 2 is that these natural datasets comprise a set of more complex environments in which learners could nonetheless still benefit from spatial generalization.

As in both previous experiments, participants (*n* = 80) sampled more locally than random chance (*t*(79) = 50.1, *p* < .001, *d* = 5.6, 95% CI (4.7,6.5), *BF* > 100), with participants in the Accumulation condition sampling more locally than those in the Maximization condition (*t*(78) = 3.1, *p* = .003, *d* = 0.7, 95% CI (0.2,1.1), *BF* = 12.1). In the natural environments, we found that Accumulation participants achieved a higher average reward than Maximization participants (*t*(78) = 2.7, *p* = .008, *d* = 0.6, 95% CI (0.2,1.1), *BF* = 5.6), with an effect size similar to Experiment 1. There was no difference in maximum reward across payoff conditions (*t*(78) = 0.3, *p* = .8, *d* = 0.06, 95% CI (–0.4,0.5), *BF* = 0.2), as in all previous experiments, showing that the goal of balancing exploration-exploitation leads to the best results on both performance metrics. As in the previous experiments, we found that a longer search horizon did not lead to higher average rewards (*t*(78) = 2.1, *p* = .04, *d* = 0.2, 95% CI (–0.2,0.7), *BF* = 0.4). The results of Experiment 3 therefore closely corroborate the results of Experiments 1 and 2, showing that findings on human behaviour in simulated environments is very similar to human behaviour in natural environments.

### Modelling Generalization and Search

To better understand how participants explore, we compared a diverse set of computational models in their ability to predict each subject’s trial-by-trial choices (see Supplementary Fig. 1 and Supplementary Table. 3 for full results). These models include different combinations of *models of learning* and *sampling strategies*, which map onto the distinction between belief and sampling models that is central to theories in statistics^35^, psychology^36^, and philosophy of science^37^. Models of learning form inductive beliefs about the value of possible options (including unobserved options) conditioned on previous observations, while sampling strategies transform these beliefs into probabilistic predictions about where a participant will sample next. We also consider heuristics, which are competitive models of human behaviour in bandit tasks^5^, yet do not maintain a model of the world (see SI). By far the best predictive models used *Gaussian Process* (GP) regression^38, 39^ as a mechanism for generalization, and *Upper Confidence Bound* (UCB) sampling^40^ as an optimistic solution to the exploration-exploitation dilemma.

Function learning provides a possible explanation of how individuals generalize from previous experience to unobserved options, by adaptively learning an underlying function mapping options onto rewards. We use GP regression as an expressive model of human function learning, which has known equivalencies to neural network function approximators^41^, yet provides psychologically interpretable parameter estimates about the extent to which generalization occurs. GP function learning can guide search by making predictions about the expected mean *m*(**x**) and the associated uncertainty *s*(**x**) (estimated here as a standard deviation) for each option x in the global state space (see Fig. 2a-b), conditioned on a finite number of previous observations of rewards **y**_*T*_ = [*y*_1_,*y*_2_,…,*y*_*T*_]^⊤^ at inputs **X**_*T*_ = [**x**_1_,…,**x**_*T*_] . Similarities between options are modelled by a *Radial Basis Function* (RBF) kernel:

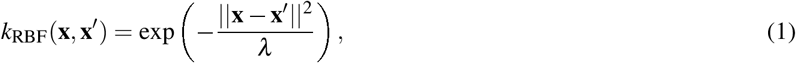

where *λ* governs how quickly correlations between points **x** and **x**̂ (e.g., two tiles on the grid) decay towards zero as their distance increases. We use *λ* as a free parameter, which can be interpreted psychologically as the extent to which people generalize spatially. Since the GP prior is completely defined by the RBF kernel, the underlying mechanisms are similar to Shepard’s universal gradient of generalization^42^, which also models generalization as an exponentially decreasing function of distance between stimuli. To illustrate, generalization to the extent of *λ* = 1 corresponds to the assumption that the rewards of two neighbouring options are correlated by *r* = 0.61, and that this correlation decays to (effectively) zero if options are further than three tiles away from each other. Smaller *λ* values would lead to a more rapid decay of assumed correlations as a function of distance.

**Figure 2.**
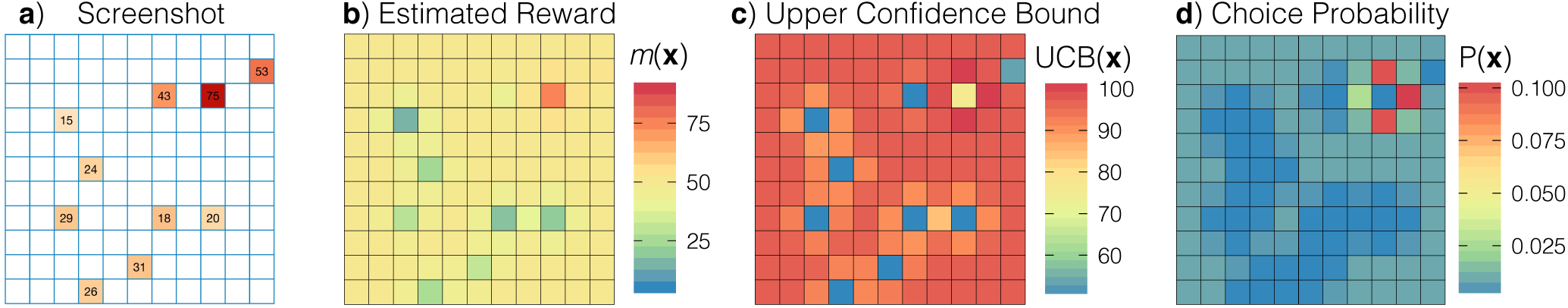
Overview of the Function Learning-UCB Model specified using median participant parameter estimates from Experiment 2 (see Supplementary Fig. 3). **a)** Screenshot of Experiment 2. Participants were allowed to select any tile until the search horizon was exhausted. **b)** Estimated reward (not shown, the estimated uncertainty) as predicted by the GP Function Learning model, based on the points sampled in Panel a. **c)** Upper confidence bound of predicted rewards. d) Choice probabilities after a softmax choice rule. 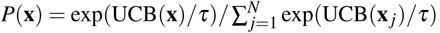, where *τ* is the temperature parameter (i.e., higher temperature values lead to more random sampling).

Given estimates about expected rewards *m*(**x**) and the underlying uncertainty *s*(**x**) from the function learning model, UCB sampling produces valuations of each option x using a simple weighted sum:

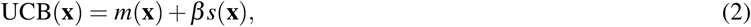

where *β* is a free parameter governing how much the reduction of uncertainty is valued relative to expectations of reward (Fig. 2c). To illustrate, an exploration bonus of *β* = 0.5 suggests participants would prefer a hypothetical option *x*_1_ predicted to have mean reward *m*(*x*_1_) = 60 and standard deviation *s*(*x*_1_) = 10, over an option *x*_2_ predicted to have mean reward *m*(*x*_2_) = 64 and standard deviation *s*(*x*_2_) = 1. This is because sampling *x*_1_ is expected to reduce a large amount of uncertainty, even though *x*_2_ has a higher mean reward (as UCB(*x*_1_) = 65 but UCB(*x*_2_) = 64.5). This trade-off between exploiting known high-value rewards and exploring to reduce uncertainty^43^ can be interpreted as optimistically inflating expectations of reward by the attached uncertainty, and can be contrasted to two separate sampling strategies that only sample based on expected reward (Pure Exploitation) or uncertainty (Pure Exploration):

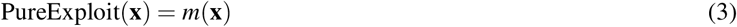

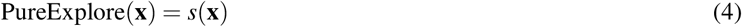

Figure 2 shows how the GP-UCB model makes inferences about the search space and uses UCB sampling (combined with a softmax choice rule) to make probabilistic predictions about where the participant will sample next. We refer to this model as the *Function Learning Model* and contrast it with an *Option Learning Model*. The Option Learning Model uses a Bayesian mean tracker to learn about the distribution of rewards for each option independently (see Methods). The Option learning Model is a traditional associative learning model, and can be understood as a variant of a Kalman filter where rewards are assumed to be time-invariant^6^. Like the Function Learning Model, the Option Learning Model also generates normally distributed predictions with mean *m*(**x**) and standard deviation *s*(**x**), which we combine with the same set of sampling strategies and the same softmax choice rule to make probabilistic predictions about search. For both models, we use the softmax temperature parameter (*τ*) to estimate the amount of undirected exploration (i.e., higher temperatures correspond to more noisy sampling; Fig. 2d), in contrast to the *β* parameter of UCB, which estimates the level of exploration directed towards reducing uncertainty.

### Modelling results

#### Experiment 1

Participants were better described by the Function Learning Model than the Option Learning Model (*t*(80) = 14.10, *p* < .001 *d* = 1.6, 95% CI (1.1,2.1), *BF* > 100, comparing cross-validated predictive accuracies, both using UCB sampling), providing evidence that participants generalized instead of learning rewards for each option independently. Furthermore, by decomposing the UCB sampling algorithm into Pure Exploit or Pure Explore components, we show that both expectations of reward and estimates of uncertainty are necessary components for the Function Learning Model to predict human search behaviour, with the Pure Exploitation (*t*(80) = –8.85, *p* < .001, *d* = –1.0, 95% CI (–0.5,−1.4), *BF* > 100) and Pure Exploration (*t*(80) = –16.63, *p* < .001, *d* = –1.8, 95% CI (–1.3,−2.4), *BF* > 100) variants each making less accurate predictions than the combined UCB algorithm.

Because of the observed tendency to sample locally, we created a localized variant of both Option Learning and Function Learning Models (indicated by an asterisk *; Fig. 3a), penalizing options farther away from the previous selected option (without introducing additional free parameters; see Methods). While the Option Learning* Model was better than the standard Option Learning Model (*t*(80) = 16.13, *p* < .001, *d* = 1.8, 95% CI (1.3,2.3), *BF* > 100), the standard Function Learning Model still outperformed its localized variant (*t*(80) = 5.05, *p* < .001, *d* = 0.6, 95% CI (0.1,1.0), *BF* > 100). Overall, 56 out of 81 participants were best described by the Function Learning Model, with an additional 10 participants best described by the Function Learning* Model with localization. Lastly, we also calculated each model’s protected probability of exceedance^44^ using its out-of-sample log-evidence. This probability assesses which model is the most common among all models in our pool (among the 12 models reported in the main text; see Supplementary Table 3 for comparison with additional models) while also correcting for chance. Doing so, we found that the Function Learning-UCB Model reached a protected probability of pxp = 1, indicating that it vastly outperformed all of the other models.

Figure 3b shows simulated learning curves of each model in comparison to human performance, where models were specified using parameters from participants’ estimates (averaged over 100 simulated experiments per participant per model). Whereas both versions of the Option Learning Model improve only very slowly, both standard and localized versions of the Function Learning Model behave sensibly and show a close alignment to the rapid rate of human learning during the early phases of learning. However, there is still a deviation in similarity between the curves, which is partially due to aggregating over reward conditions and horizon manipulations, in addition to aggregating over individuals, where some participants over-explore their environments while others produce continuously increasing learning curves (see Supplementary Figure 6 for individual learning curves). While aggregated learning curves should be analyzed with caution^45^, we find an overlap between elements of human intelligence responsible for successful performance in our task, and elements of participant behaviour captured by the Function Learning Model.

We compare participants’ parameter estimates using a Wilcoxon signed rank test to make the resulting differences more robust to potential outliers. The parameter estimates of the Function Learning Model (Fig. 3c) indicated that people tend to underestimate the extent of spatial correlations, with median per-participant *λ* estimates significantly lower than the ground truth (*λ*_*Smooth*_ = 2 and *λ*_*Rough*_ = 1) for both Smooth (Wilcoxon signed rank test; *λ̃*_*Smooth*_ = 0.5, *Z* = –7.1, *p* < .001, *r* = 1.1, *BF*_*Z*_ > 100) and Rough environments (*λ̃*_*Rough*_ = 0.5, *Z* = –3.4, *p* < .001, *r* = 0.55, *BF*_*Z*_ > 100). This can be interpreted as a tendency towards undergeneralization. Additionally, we found that the estimated exploration bonus of UCB sampling (*b*) was reliably greater than zero (*β̃* = 0.51, *Z* = –7.7, *p* < .001, *r* = 0.86, *BF*_*Z*_ > 100, compared to lower estimation bound), reflecting the valuation of sampling uncertain options, together with exploiting high expectations of reward. Lastly, we found relatively low estimates of the softmax temperature parameter (*τ̃* = 0.01), suggesting that the search behaviour of participants corresponded closely to selecting the very best option, once they had taken into account both the exploitation and exploration components of the available actions.

**Figure 3.**
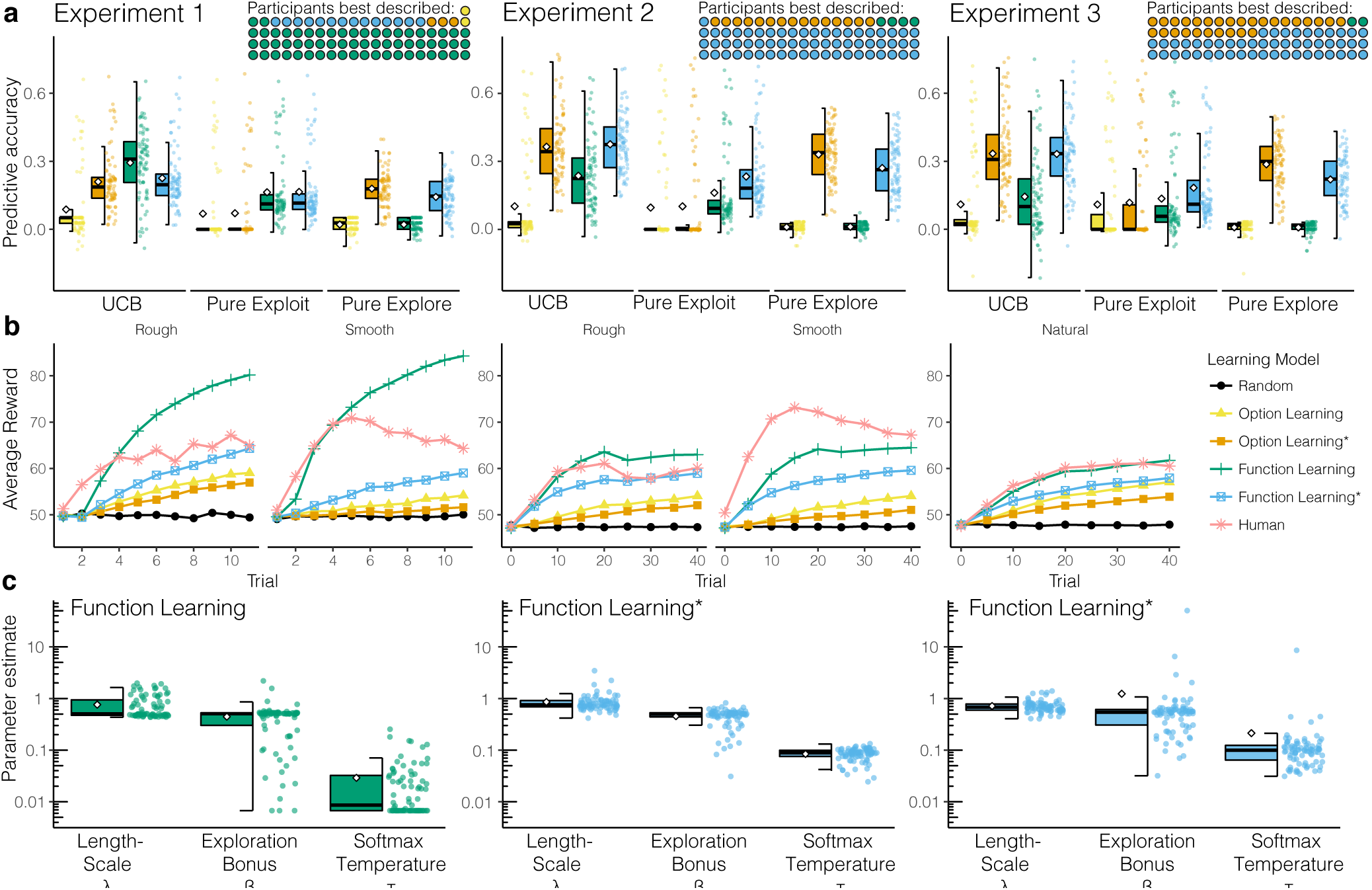
Modelling results. **a)** Cross-validated predictive accuracy of each model (higher is better), with box plots indicating the IQR, the median (horizontal line), mean (diamond), and 1.5x IQR (whiskers). Each individual participant is shown as a single dot, with the number of participants best described shown as an icon array (inset; aggregated by sampling strategies). Asterisks (*) indicate a localized variant of the Option Learning or Function Learning models, where predictions are weighted by the inverse distance from the previous choice (see Methods). **b)** Learning curves of participants and model simulations. Each simulated learning model uses UCB sampling and is specified using participants parameter estimates and averaged over 100 simulated experiments per participant per model. **c)** Parameter estimates of the best predicting model for each experiment. Each coloured dot is the median estimate per participant, with box plots indicating 1.5x IQR (whiskers), median (horizontal line), and mean (diamond).

#### Experiment 2

In a more complex bivariate environment (Fig. 3a), the Function Learning Model again made better predictions than the Option Learning Model (*t*(79) = 9.99, *p* < .001, *d* = 1.1, 95% CI (0.6,1.6), *BF* > 100), although this was only marginally the case when comparing localized Function Learning* to localized Option Learning* (*t*(79) = 2.05, *p* = .044, *d* = 0.2, 95% CI (–0.2,0.7), *BF* = 0.9). In the two-dimensional search environment of Experiment 2, adding localization improved predictions for both Option Learning (*t*(79) = 19.92, *p* < .001, *d* = 2.2, 95% CI (1.7,2.8), *BF* > 100) and Function Learning (*t*(79) = 10.47, *p* < .001, *d* = 1.2, 95% CI (0.7,1.6), *BF* > 100), in line with the stronger tendency towards localized sampling compared to Experiment 1 (see Fig. 1c). Altogether, 61 out of 80 participants were best predicted by the localized Function Learning* model, whereas only 12 participants were best predicted by the localized Option Learning* model. Again, both components of the UCB strategy were necessary to predict choices, with Pure Exploit (*t*(79) = –6.44, *p* < .001, *d* = –0.7, 95% CI (–0.3,−1.2), *BF* > 100) and Pure Explore (*t*(79) = –12.8, *p* < .001, *d* = –1.4, 95% CI (–0.9,−1.9), *BF* > 100) making worse predictions. The probability of exceedance over all models showed that the Function Learning*-UCB Model achieved virtually pxp = 1, indicating that it greatly outperformed all other models under consideration.

As in Experiment 1, the simulated learning curves of the Option Learning models learned slowly and only marginally outperformed a random sampling strategy (Fig. 3b), whereas both variants of the Function Learning Model achieved performance comparable to that of human participants. Median per-participant parameter estimates (Fig. 3c) from the Function Learning*-UCB Model showed that while participants generalized somewhat more than in Experiment 1 (*λ̂* = 0.75, *Z* = –3.7, *p* < .001, *r* = 0.29, *BF*_*Z*_ > 100), they again underestimated the strength of the underlying spatial correlation in both Smooth (*λ̂*_*Smooth*_ = 0.78, *Z* = –5.8, *p* < .001, *r* = 0.88, *BF*_*Z*_ > 100; comparison to *λ*_*Smooth*_ = 2) and Rough environments (*λ̃*_*Rough*_ = 0.75, *Z* = –4.7, *p* < .001, *r* = 0.78; comparison to *λ*_*Rough*_ = 1, *BF*_*Z*_ > 100). This suggests a robust tendency to undergeneralize. There were no differences in the estimated exploration bonus *β* between Experiment 1 and 2 (*β̃* = 0.5, *Z* = 0.86, *p* = .80, *r* = 0.07, *BF*_*Z*_ = 0.2), although the estimated softmax temperature parameter *τ* was larger than in Experiment 1 (*τ̃* = 0.09; *Z* = –8.89, *p* < .001, *r* = 0.70, *BF*_*Z*_ = 34). Experiment 2 therefore replicated the main findings of Experiment 1. When taken together, results from the two experiments provide strong evidence that human search behaviour is best explained by function learning paired with an optimistic trade-off between exploration and exploitation.

#### Experiment 3

Using natural environments without a fixed level of spatial correlations, we replicated key results from the prior experiments: Function Learning made better predictions than Option Learning (*t*(79) = 3.03, *p* = .003, *d* = 0.3, 95% CI (–0.1,0.8), *BF* = 8.2); adding localization improved predictions for both Option Learning (*t*(79) = 18.83, *p* < .001, *d* = 2.1, 95% CI (1.6,2.6), *BF* > 100) and Function Learning (*t*(79) = 14.61, *p* < .001, *d* = 1.6, 95% CI (1.1,2.1), *BF* > 100); and the combined UCB algorithm performed better than using only a Pure Exploit (*t*(79) = 12.97, *p* < .001, *d* = 1.4, 95% CI (1.0,1.9), *BF* > 100) or a Pure Explore strategy (*t*(79) = 5.87, *p* < .001, *d* = 0.7, 95% CI (0.3,1.2), *BF* > 100). However, the difference between the localized Function Learning* and the localized Option Learning* was negligible (*t*(79) = 0.32, *p* = .75, *d* = 0.04, 95% CI (–0.4,0.5), *BF* = 0.1). This is perhaps due to the high variability across environments, which makes it harder to predict out-of-sample choices using generalization behaviour (i.e., *λ*) estimated from a separate set of environments. Nevertheless, the localized Function Learning* model was still the best predicting model for the majority of participants (48 out of 80 participants). Moreover, calculating the protected probability of exceedance over all models’ predictive evidence revealed a probability of pxp = 0.98 that the Function Learning* model was more frequent in the population than all the other models, followed by pxp = 0.01 for the Option Learning* model. Thus, even in natural environments in which the underlying spatial correlations are unknown, we were still able to distinguish the different models in terms of their overall out-of-sample predictive performance.

The simulated learning curves in Figure 3b show the strongest concurrence out of all previous experiments between the Function Learning model and human performance. Moreover, both variants of the Option Learning model learn far slower, failing to match the rate of human learning, suggesting that they are not plausible models of human behaviour^46^. The parameter estimates from the Function Learning* Model are largely consistent with the results from Experiment 2 (Fig. 3c), but with participants generalizing slightly less (*λ̂*_*natural*_ = 0.68, *Z* = –3.4, *p* < .001, *r* = 0.27, *BF*_*Z*_ = 9.6), and exploring slightly more, with a small increase in both directed exploration (*β̂*_*natural*_ = 0.54, *Z* = –2.3, *p* = .01, *r* = 0.18, *BF*_*Z*_ = 4.5) and undirected exploration (*τ̂*_*natural*_ = 0.1, *Z* = –2.2, *p* = .02, *r* = 0.17, *BF*_*Z*_ = 4.2) parameters. Altogether, the parameter estimates are highly similar to the previous experiments.

### Robustness and Recovery

We conducted both model and parameter recovery simulations to assess the validity of our modelling results (see SI). Model recovery consisted of simulating data using a *generating model* specified by participant parameter estimates. We then performed the same cross-validation procedure to fit a *recovering model* on this simulated data. In all cases, the best predictive accuracy occurred when the recovering model matched the generating model (Supplementary Figure 2), suggesting robustness to Type I errors and ruling out model overfitting (i.e., the Function Learning Model did not best predict data generated by the Option Learning Model). Parameter recovery was performed to ensure that each parameter in the Function Learning-UCB Model robustly captured separate and distinct phenomena. In all cases, the generating and recovered parameter estimates were highly correlated (Supplementary Figure 3). It is noteworthy that we found distinct and recoverable estimates for *β* (exploration bonus) and *τ* (softmax temperature), supporting the existence of exploration *directed* towards reducing uncertainty^12^ as a separate phenomena from noisy, undirected exploration^47^.

### The Adaptive Nature of Undergeneralization

In Experiments 1 and 2, we observed a robust tendency to undergeneralize compared to the true level of spatial correlations in the environment. We therefore ran simulations to assess how different levels of generalization influence search performance when paired with different types of environments. We found that undergeneralization largely leads to better performance than overgeneralization. Remarkably, undergeneralization sometimes is even better than exactly matching the underlying structure of the environment (Fig. 4). These simulations were performed by first generating search environments by sampling from a GP prior specified using a *teacher* length-scale (*λ*_0_), and then simulating search in this environment by specifying the Function Learning-UCB Model with a *student* length-scale (*λ*_1_). Instead of a discrete grid, we chose a set-up common in Bayesian optimization^48^ with continuous bivariate inputs in the range *x*,*y* = [0,1], allowing for a broader set of potential mismatched alignments (see Supplemental Figure 4 for simulations using the exact design of each experiment).

**Figure 4.**
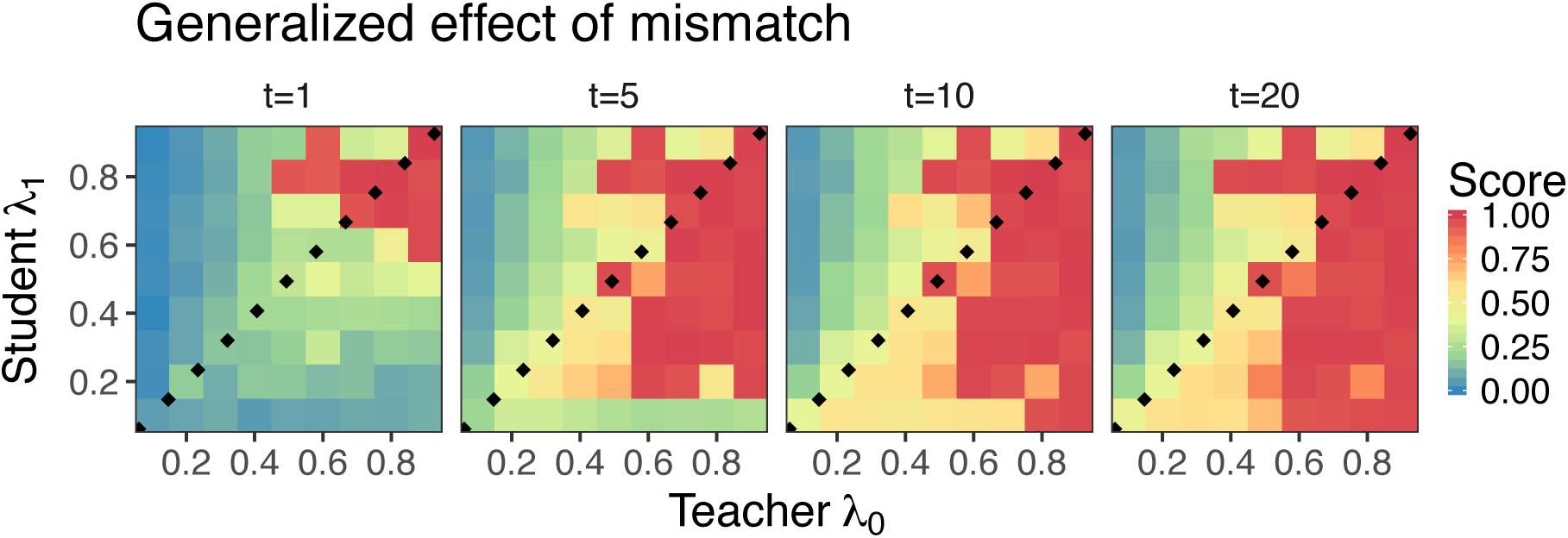
Mismatched length-scale (*λ*) simulation results. The teacher length-scale *λ*_0_ is on the x-axis, the student length-scale *λ*_1_ is on the y-axis, and each panel is performance at a different trial *t*. The teacher *λ*_0_ values were used to generate environments, while the student *λ*_1_ values were used to parameterize the Function Learning Model to simulate search performance. The dotted lines show where *λ*_0_ = *λ*_1_ and mark the difference between undergeneralization and overgeneralization, with points below the line indicating undergeneralization. We report the median score (over 100 replications) as a standardized measure of performance, such that 0 shows the lowest possible and 1 the highest possible log unit-performance.

We find that undergeneralization largely leads to better performance than overgeneralization, and that this effect is more pronounced over time *t* (i.e., longer search horizons). Estimating the best possible alignment between *λ*_0_ and *λ*_1_ revealed that underestimating *λ*_0_ by an average of about 0.21 produces the best scores over all scenarios. These simulation results show that the systematically lower estimates of *λ* captured by our models are not necessarily a flaw in human cognition, but can sometimes lead to better performance. Indeed, simulations based on the natural environments used in Experiment 3 (which had no fixed level of spatial correlations) revealed that the range of participant *λ* estimates were highly adaptive to the environments they encountered (Supplemental Figure 4c). Undergeneralization might not be a bug, but rather an important feature of human behaviour.

## Discussion

How do people learn and adaptively make good decisions when the number of possible actions is vast and not all possibilities can be explored? We found that Function Learning, operationalized using GP regression, provides a mechanism for generalization, which can be used to guide search towards unexplored yet promising options. Combined with Upper Confidence Bound (UCB) sampling, this model navigates the exploration-exploitation dilemma by optimistically inflating expectations of reward by the estimated uncertainty.

While GP function learning combined with a UCB sampling algorithm has been successfully applied to search problems in ecology^49^, robotics^50, 51^, and biology^52^, there has been little psychological research on how humans learn and search in environments with a vast set of possible actions. The question of how generalization operates in an active learning context is of great importance, and our work makes key theoretical and empirical contributions. Expanding on previous studies that found an overlap between GP-UCB and human learning rates^8, 23^, we use cognitive modelling to understand how humans generalize and address the exploration-exploitation dilemma in a complex search task with spatially correlated outcomes.

Through multiple analyses, including trial-by-trial predictive cross-validation and simulated behaviour using participants’ parameter estimates, we competitively assessed which models best predicted human behaviour. The vast majority of participants were best described by the Function Learning-UCB model or its localized variant. Parameter estimates from the best-fitting Function Learning-UCB models suggest there was a systematic tendency to undergeneralize the extent of spatial correlations, which we found can sometimes lead to better search performance than even an exact match with the underlying structure of the environment (Fig. 4).

Altogether, our modelling framework yielded highly robust and recoverable results (Supplemental Figure 2) and parameter estimates (Supplemental Figure 3). Whereas previous research on exploration bonuses has had mixed results^6, 12, 47^, we found recoverable parameter estimates for the separate phenomena of directed exploration, encoded in UCB exploration parameter *β*, and the noisy, undirected exploration encoded in the softmax temperature parameter *τ*. Even though UCB sampling is both *optimistic* (always treating uncertainty as positive) and *myopic* (only planning the next timestep), similar algorithms have competitive performance guarantees in a bandit setting^53^. This shows a remarkable concurrence between intuitive human strategies and state-of-the-art machine learning research.

### Limitations and extensions

One potential limitation is that our payoff manipulation failed to induce superior performance according to the relevant performance metric. While participants in the Accumulation condition achieved higher average reward, participants in the Maximization condition were not able to outperform with respect to the maximum reward criterion. The goal of balancing exploration-exploitation (Accumulation condition) or the goal of global optimization (Maximization condition) was induced through the manipulation of written instructions, comprehension check questions, and feedback between rounds (see Methods). While this may have been insufficient for observing clear performance differences (see SI for parameter differences), the practical difference between these two goals is murky even in the Bayesian optimization literature, where the strict goal of finding the global optimum is often abandoned based purely on computational concerns^54^. Instead, the global optimization goal is frequently replaced by an approximate measure of performance, such as cumulative regret^53^, which closely aligns to our Accumulation payoff condition. In our experiments, remarkably, participants assigned to the Accumulation goal payoff condition also performed best relative to the maximization criterion.

In addition to providing the best model of human behaviour, the Function Learning Model also offers many opportunities for theory integration. The Option Learning Model can itself be reformulated as special case of GP regression^55^. When the length-scale of the RBF kernel approaches zero (*λ* → 0), the Function Learning Model assumes state independence, as in the Option Learning Model. Thus, there may be a continuum of reinforcement learning models, ranging from the traditional assumption of state *in*dependence to the opposite extreme, of complete state *inter*-dependence. Moreover, GPs also have equivalencies to Bayesian neural networks^41^, suggesting a further link to distributed function learning models^56^. Indeed, one explanation for the impressive performance of deep reinforcement learning^14^ is that neural networks are specifically a powerful type of function approximator^57^.

Lastly, both spatial and conceptual representations have been connected to a common neural substrate in the hippocampus^27^, suggesting a potential avenue for applying the same Function Learning-UCB model for modelling human learning using contextual^28–30^, semantic^31, 32^, or potentially even graph-based features. One hypothesis for this common role of the hippocamus is that it performs predictive coding of future state transitions^58^, also known as “successor representation”^24^. In our task, where there are no restrictions on state transitions (i.e., each state is reachable from any prior state), it may be the case that the RBF kernel driving our GP Function Learning model performs the same role as the transition matrix of a successor representation model, where state transitions are learned via a random walk policy.

## Conclusions

We present a paradigm for studying how people use generalization to guide the active search for rewards, and found a systematic—yet sometimes beneficial—tendency to undergeneralize. Additionally, we uncovered substantial evidence for the separate phenomena of directed exploration (towards reducing uncertainty) and noisy, undirected exploration. Even though our current implementation only grazes the surface of the types of complex tasks people are able to solve—and indeed could be extended in future studies using temporal dynamics or depleting resources—it is far richer in both the set-up and modelling framework than traditional multi-armed bandit problems used for studying human behaviour. Our empirical and modelling results show how function learning, combined with optimistic search strategies, may provide the foundation of adaptive behaviour in complex environments.

## Methods

*Participants* 81 participants were recruited from Amazon Mechanical Turk for Experiment 1 (25 Female; mean ± SD age 33 ± 11), 80 for Experiment 2 (25 Female; mean ± SD age 32 ± 9), and 80 for Experiment 3 (24 Female; mean ± SD age 35 ± 10). In all experiments, participants were paid a participation fee of $0.50 and a performance contingent bonus of up to $1.50. Participants earned on average $1.14 ± 0.13 and spent 8 ± 4 minutes on the task in Experiment 1, earned $1.64 ± 0.20 and spent 8 ± 4 minutes in Experiment 2, and earned $1.53 ± 0.15 and spent 8 ± 5 minutes in Experiment 3. Participants were only allowed to participate in one of the experiments, and were required to have a 95% HIT approval rate and 1000 previously completed HITs. No statistical methods were used to pre-determine sample sizes but our sample sizes are similar or larger to those reported in previous publications^6, 12, 23, 28, 29^. The Ethics Committee of the Max Planck Institute for Human Development approved the methodology and all participants consented to participation through an online consent form at the beginning of the survey.

### Design

Experiments 1 and 2 used a 2×2 between-subjects design, where participants were randomly assigned to one of two different payoff structures (*Accumulation condition* vs. *Maximization condition*) and one of two different classes of environments (*Smooth* vs. *Rough*), whereas Experiment 3 used environments from real-world agricultural datasets, and manipulated only the payoff structure (random assignment between subjects). Each grid world represented a (either uni-or bivariate) function, with each observation including normally distributed noise, *ε* ~ *𝒩* (0,1). The task was presented over either 16 rounds (Exp.1) or 8 rounds (Exp. 2 and 3) on different grid worlds, which were randomly drawn (without replacement) from the same class of environments. Participants had either a short or long search horizon (Exp. 1: [5,10]; Exp. 2 and 3: [20, 40]) to sample tiles on the grid, including repeat clicks. The search horizon alternated between rounds (within subject), with initial horizon length counterbalanced between subjects by random assignment. Data collection and analysis were not performed blind to the conditions of the experiments.

### Materials and procedure

Prior to starting the task, participants observed four fully revealed example environments and had to correctly complete three comprehension questions. At the beginning of each round, one random tile was revealed and participants could click any of the tiles in the grid until the search horizon was exhausted, including re-clicking previously revealed tiles. Clicking an unrevealed tile displayed the numerical value of the reward along with a corresponding colour aid, where darker colours indicated higher point values. Per round, observations were scaled to a randomly drawn maximum value in the range of 65 to 85, so that the value of the global optima could not be easily guessed (e.g., a value of 100). Re-clicked tiles could show some variations in the observed value due to noise. For repeat clicks, the most recent observation was displayed numerically, while hovering over the tile would display the entire history of observation. The colour of the tile corresponded to the mean of all previous observations.

#### Payoff conditions

We compared performance under two different payoff conditions, requiring either a balance between exploration and exploitation (*Accumulation condition*) or corresponding to consistently making exploration decisions (*Maximization condition*). In each payoff condition, participants received a performance contingent bonus of up to $1.50. *Accumulation condition* participants were given a bonus based on the average value of all clicks as a fraction of the global optima, 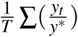, where *y*^*^ is the global optimum, whereas participants in the *Maximization condition* were rewarded using the ratio of the highest observed reward to the global optimum, 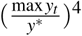, taken to the power of 4 to exaggerate differences in the upper range of performance and for between-group parity in expected earnings across payoff conditions. Both conditions were equally weighted across all rounds and used noisy but unscaled observations to assign a bonus of up to $1.50. Subjects were informed in dollars about the bonus earned at the end of each round.

#### Environments

In Experiments 1 and 2, we used two classes of generated environments corresponding to different levels of smoothness (i.e., spatial correlation of rewards). These environments were sampled from a GP prior with a RBF kernel, where the length-scale parameter (*λ*) determines the rate at which the correlations of rewards decay over distance. *Rough* environments used *λ*_*Rough*_ = 1 and *Smooth* environments used *λ*_*Smooth*_ = 2, with 40 environments (Exp. 1) and 20 environments (Exp. 2) generated for each class (Smooth and Rough). In Experiment 3, we used environments defined by 20 real-world agricultural datasets, where the location on the grid corresponds to the rows and columns of a field and the payoffs reflect the normalized yield of various crops (see SI for full details).

#### Search horizons

We chose two horizon lengths (Short=5 or 20 and Long=10 or 40) that were fewer than the total number of tiles on the grid (30 or 121), and varied them within subject (alternating between rounds and counterbalanced). Horizon length was approximately equivalent between Experiments 1 and Experiments 2 and 3, as a fraction of the total number of options 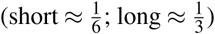.

### Statistical tests

All reported *t*-tests are two-sided. We also report Bayes Factors quantifying the likelihood of the data under *H*_*A*_ relative to the likelihood of the data under *H*_0_. We calculate the default two-sided Bayesian *t*-test using a Jeffreys-Zellner-Siow prior with its scale set to 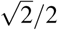, following^59^. For parametric tests, the data distribution was assumed to be normal but this was not formally tested. For non-parametric comparisons, the Bayes Factor *BF*_*Z*_ is derived by performing posterior inference over the Wilcoxon test statistics and assigning a prior by means of a parametric yoking procedure^60^. The null hypothesis posits that the statistic between two groups does not differ and the alternative hypothesis posits the presence of an effect and assigns an effect size using a Cauchy distribution with the scale parameter set to 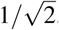.

### Localization of Models

To penalize search options by the distance from the previous choice, we weighted each option by the inverse Manhattan distance (IMD) to the last revealed tile 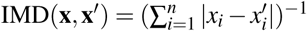, prior to the softmax transformation. For the special case where **x** = **x**′, we set IMD(**x**,**x**′) = 1. Localized models are indicated by an asterix (*).

### Model Comparison

We performed model comparison using cross-validated maximum likelihood estimation (MLE), where each participant’s data was separated by horizon length (short or long) and we iteratively form a training set by leaving out a single round, compute a MLE on the training set, and then generate out-of-sample predictions on the remaining round (see SI for further details). This was repeated for all combinations of training set and test set, and for both short and long horizons. The cross-validation procedure yielded one set of parameter estimates per round, per participant, and out-of-sample predictions for 120 choices in Experiment 1 and 240 choices in Experiments 2 and 3 (per participant). Prediction error (computed as log loss) was summed up over all rounds, and is reported as *predictive accuracy*, using a pseudo-*R*^2^ measure that compares the total log loss prediction error for each model to that of a random model:

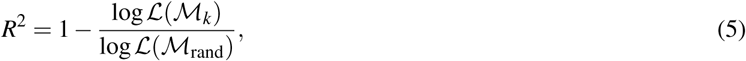

where log*𝓛*(*𝓜*_rand_) is the log loss of a random model and log*𝓛*(*𝓜*_*k*_) is model *k*’s out-of-sample prediction error. Moreover, we calculated each model’s protected probability of exceedance using its predictive log-evidence^44^. This probability is defined as the probability that a particular model is more frequent in the population than all the other models, averaged over the probability of the null hypothesis that all models are equally frequent (thereby correcting for chance performance).

## Code Availability

Code for all models and analyses is available at https://github.com/charleywu/gridsearch

## Data Availability

Anonymised participant data and model simulation data are available at https://github.com/charleywu/gridsearch

## Author Contributions

C.M.W. and E.S. designed the experiments, collected and analysed the data, and wrote the paper. M.S., J.D.N., and B.M. designed the experiments and wrote the paper.

## Acknowledgments

We thank Peter Todd, Tim Pleskac, Neil Bramley, Henrik Singmann, and Mehdi Moussaïd for helpful feedback. This work was supported by the International Max Planck Research School on Adapting Behavior in a Fundamentally Uncertain World (CMW), by the Harvard Data Science Initiative (ES), and DFG grants ME 3717/2-2 to BM and NE 1713/1-2 to JDN as part of the New Frameworks of Rationality (SPP 1516) priority program. The funders had no role in study design, data collection and analysis, decision to publish or preparation of the manuscript.

## Supporting information

### Full Model Comparison

We report the full model comparison of 27 models, of which 12 (i.e., four learning models and three sampling strategies) are included in the main text. We use different *Models of Learning* (i.e., Function Learning and Option Learning), which combined with a *Sampling Strategy* can make predictions about where a participant will search, given the history of previous observations. We also include comparisons to *Simple Heuristic Strategies*^61^, which make predictions about search decisions without maintaining a representation of the world (i.e., without a learning model). Table S3 shows the predictive accuracy, the number of participants best described, the protected probability of exceedance and the median parameter estimates of each model. Figure S1 shows a more detailed assessment of predictive accuracy and model performance, with participants separated by payoff condition and environment type.

**Figure S1.**
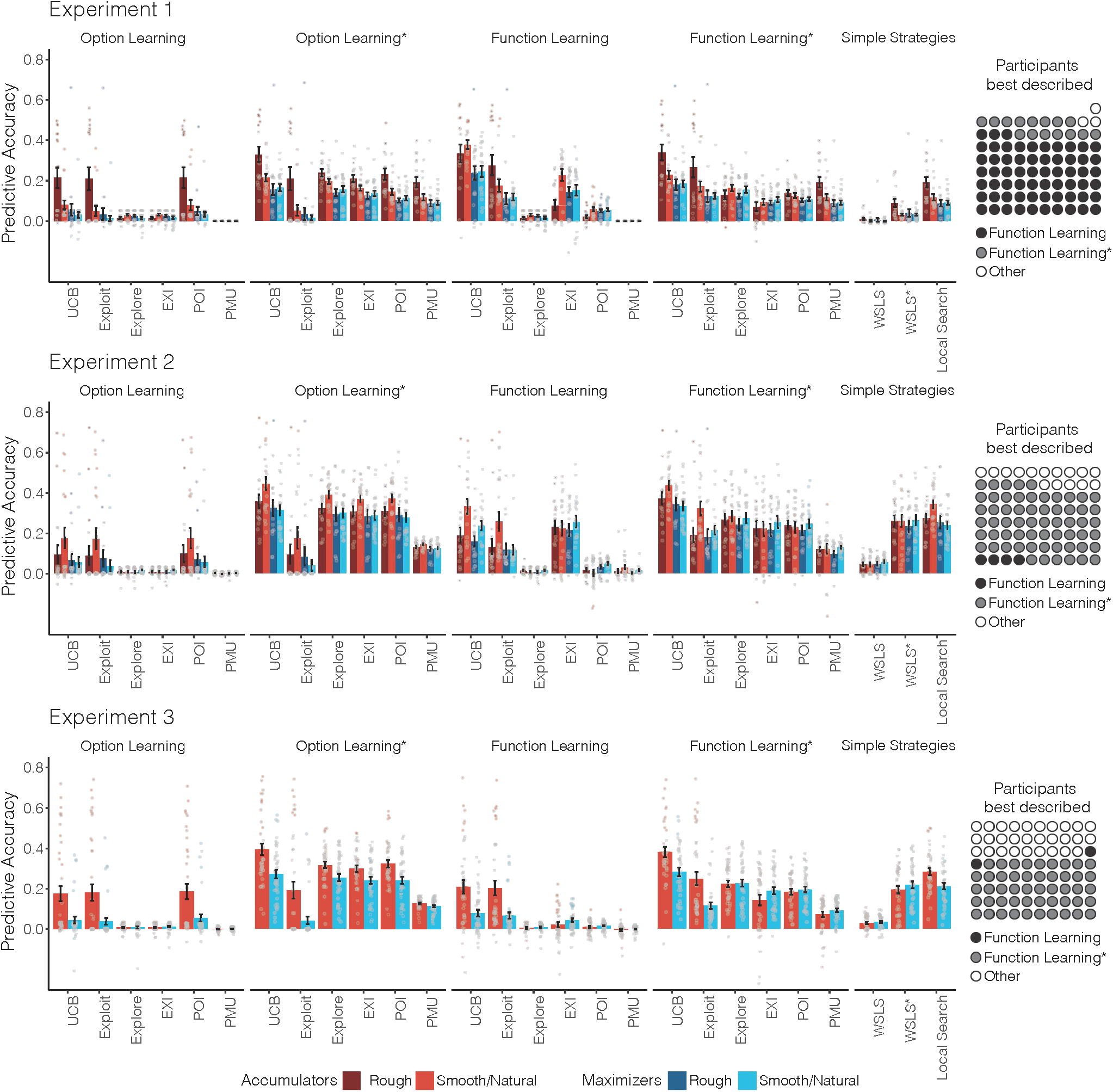
Full model comparison of all 27 models. The learning model is indicated above (or lack of in the case of simple heuristic strategies), and sampling strategy are along the x-axis. Bars indicate predictive accuracy (group mean) along with standard error, and are separated by payoff condition (colour) and environment type (darkness), with individual participants overlaid as dots. Icon arrays (right) show the number participants best described (out of the full 27 models) and are aggregated over payoff conditions, environment types, and sampling strategy. Table S3 provides more detail about the number of participants best described by each model as well as the protected probability of exceedance.

### Models of Learning

#### Function Learning.

The Function Learning Model adaptively learns an underlying function mapping spatial locations onto rewards. We use Gaussian Process (GP) regression as a Bayesian method of function learning^39^. A GP is defined as a collection of points, any subset of which is multivariate Gaussian. Let *f* : *𝒳* → R^*n*^ denote a function over input space *𝒳* that maps to real-valued scalar outputs. This function can be modelled as a random draw from a GP:

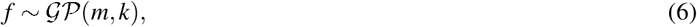

where *m* is a mean function specifying the expected output of the function given input x, and *k* is a kernel (or covariance) function specifying the covariance between outputs.

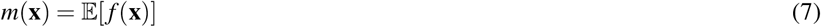

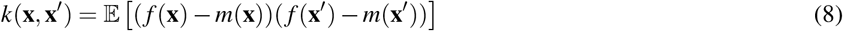

Here, we fix the prior mean to the median value of payoffs, *m*(**x**) = 50 and use the kernel function to encode an inductive bias about the expected spatial correlations between rewards (see Radial Basis Function kernel). Conditional on observed data 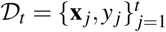, where *y*_*j*_ ~ *𝒩* (*f* (**x**_*j*_),*σ*^2^) is drawn from the underlying function with added noise *σ*^2^ = 1, we can calculate the posterior predictive distribution for a new input **x**_*_ as a Gaussian with mean *m*_*t*_(**x**_*_) and variance *v*_*t*_(**x**_*_) given by:

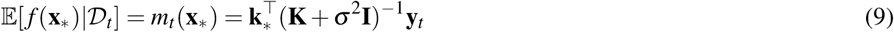

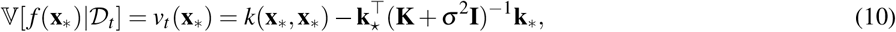

where **y** = [*y*_1_,…,*y*_*t*_]^⊤^, **K** is the *t* × *t* covariance matrix evaluated at each pair of observed inputs, and **k**_*_ = [*k*(**x**_1_,**x**_*_),…,*k*(**x**_*t*_,**x**_*_)] is the covariance between each observed input and the new input **x**_*_.

We use the Radial Basis Function (RBF) kernel as a component of the GP function learning algorithm, which specifies the correlation between inputs.

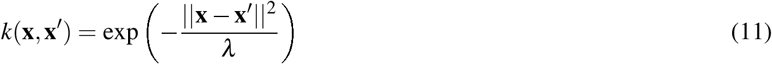

This kernel defines a universal function learning engine based on the principles of Bayesian regression and can model any stationary function. Note, sometimes the RBF kernel is specified as 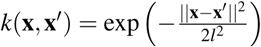 whereas we use *λ* = 2*l*^2^ as a more psychologically interpretable formulation. Intuitively, the RBF kernel models the correlation between points as an exponentially decreasing function of their distance. Here, *λ* modifies the rate of correlation decay, with larger *λ*-values corresponding to slower decays, stronger spatial correlations, and smoother functions. As *λ* → +∞, the RBF kernel assumes functions approaching linearity, whereas as *λ* → 0, there ceases to be any spatial correlation, with the implication that learning happens independently for each input without generalization (similar to traditional models of associative learning). We treat *λ* as a free parameter, and use cross-validated estimates to make inferences about the extent to which participants generalize.

#### Option Learning.

The Option Learning Model uses a Bayesian Mean Tracker, which is a type of associative learning model that assumes the average reward associated with each option is constant over time (i.e., no temporal dynamics, as opposed to the assumptions of a Kalman filter or Temporal Difference Learning)^6^, as is the case in our experimental search tasks. In contrast to the Function Learning model, the Option Learning model learns the rewards of each option separately, by computing an independent posterior distribution for the mean *μ*_*j*_ for each option *j*. We implement a version that assumes rewards are normally distributed (as in the GP Function Learning Model), with a known variance but unknown mean, where the prior distribution of the mean is again a normal distribution. This implies that the posterior distribution for each mean is also a normal distribution:

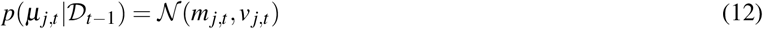

For a given option *j*, the posterior mean *m*_*j*,*t*_ and variance *v*_*j*,*t*_ are only updated when it has been selected at trial *t*:

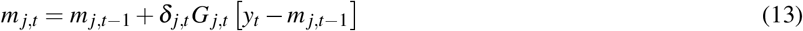

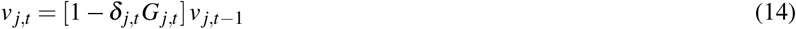

where *δ*_*j*,*t*_ = 1 if option *j* was chosen on trial *t*, and 0 otherwise. Additionally, *y*_*t*_ is the observed reward at trial *t*, and *G*_*j*,*t*_ is defined as:

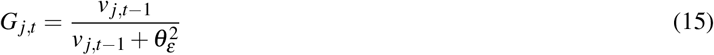

where 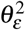 is the error variance, which is estimated as a free parameter. Intuitively, the estimated mean of the chosen option *m*_*j*,*t*_ is updated based on the difference between the observed value *y*_*t*_ and the prior expected mean *m*_*j*,*t*–1_, multiplied by *G*_*j*,*t*_. At the same time, the estimated variance *v*_*j*,*t*_ is reduced by a factor of 1 – *G*_*j*,*t*_, which is in the range [0,1]. The error variance (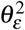) can be interpreted as an inverse sensitivity, where smaller values result in more substantial updates to the mean *m*_*j*,*t*_, and larger reductions of uncertainty *v*_*j*,*t*_. We set the prior mean to the median value of payoffs *m*_*j*,0_ = 50 and the prior variance *v*_*j*,0_ = 500.

### Sampling Strategies

Given the normally distributed posteriors of the expected rewards, which have mean *m*_*t*_(**x**) and the estimated uncertainty (estimated here as a standard deviation) 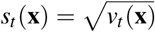, for each search option **x** (for the Option Learning model, we let *m*_*t*_(**x**) = *m*_*j*,*t*_ and *v*_*t*_(**x**) = *v*_*j*,*t*_, where *j* is the index of the option characterized by **x**), we assess different sampling strategies that (with a softmax choice rule) make probabilistic predictions about where participants search next at time *t* + 1.

#### Upper Confidence Bound Sampling.

Given the posterior predictive mean *m*_*t*_(**x**) and the estimated uncertainty *s*_*t*_(**x**), we calculate the upper confidence bound (UCB) using a simple weighted sum

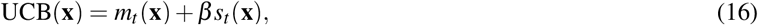

where the exploration factor *β* determines how much reduction of uncertainty is valued (relative to exploiting known high-value options) and is estimated as a free parameter.

#### Pure Exploitation and Pure Exploration.

Upper Confidence Bound sampling can be decomposed into a Pure Exploitation component, which only samples options with high expected rewards, and a Pure Exploration component, which only samples options with high uncertainty.

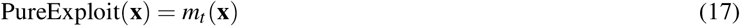

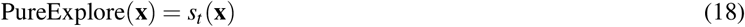

#### Expected Improvement.

At any point in time *t*, the best observed outcome can be described as **x**^+^ = argmax_**x**_*i*_∈**x**_1:*t*__ *m*_*t*_(**x**_*i*_). Expected Improvement (EXI) evaluates each option by *how much* (in the expectation) it promises to be better than the best observed outcome **x**^+^:

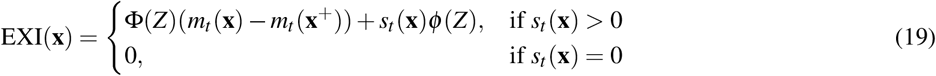

where Φ(·) is the normal CDF, *ϕ*(·) is the normal PDF, and *Z* = (*m*_*t*_(**x**) – *m*_*t*_(**x**^+^))/*s*_*t*_(**x**).

#### Probability of Improvement.

The Probability of Improvement (POI) strategy evaluates an option based on *how likely* it will be better than the best outcome (**x**^+^) observed so far:

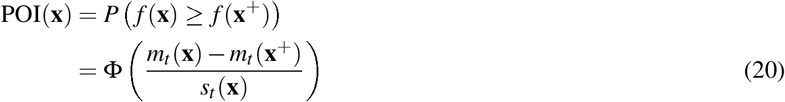

#### Probability of Maximum Utility.

The Probability of Maximum Utility (PMU) samples each option according to the probability that it results in the highest reward of all options in a particular context^6^. It is a form of probability matching and can be implemented by sampling from each option’s predictive distributions, and then choosing each option proportional to the number of times it has the highest sampled payoff.

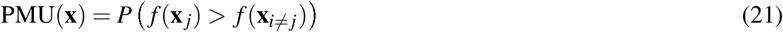

We implement this sampling strategy by Monte Carlo sampling from the posterior predictive distribution of a learning model for each option, and evaluating how often a given option turns out to be the maximum over 1,000 generated samples.

### Simple Heuristic Strategies

We also compare various simple heuristic strategies that make predictions about search behaviour without learning about the distribution of rewards.

#### Win-Stay Lose-Sample.

We consider a form of a win-stay lose-sample (WSLS) heuristic^62^, where a *win* is defined as finding a payoff with a higher or equal value than the previously best observed outcome. When the decision-maker “wins”, we assume that any tile with a Manhattan distance ≤ 1 is chosen (i.e., a repeat or any of the four cardinal neighbours) with equal probability. *Losing* is defined as the failure to improve, and results in sampling any unrevealed tile with equal probability.

#### Local Search.

Local search predicts that search decisions have a tendency to stay local to the previous choice. We use inverse Manhattan distance (IMD) to quantify locality:

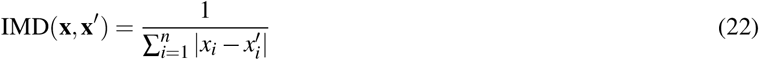

where **x** and **x**′ are vectors in R^*n*^. For the special case where **x** = **x**′, we set IMD(**x**,**x**′) = 1.

### Localization of Models

With the exception of the *Local Search* model, all other models include a localized variant, which introduced a locality bias by weighting the predicted value of each option *q*(**x**) by the inverse Manhattan distance (IMD) to the previously revealed tile. This is equivalent to a multiplicative combination with the Local Search model, similar to a “stickiness parameter”^63, 64^, although we implement it here without the introduction of any additional free parameters. Localized models are indicated with an asterisk (e.g., Function Learning*).

### Model Comparison

We use maximum likelihood estimation (MLE) for parameter estimation, and cross-validation to measure out-of-sample predictive accuracy as well as the probability of exceedance to estimate a model’s posterior probability to be the underlying predictive model of our task, given the pool of all models in our comparison. A softmax choice rule transforms each model’s valuations into a probability distribution over options:

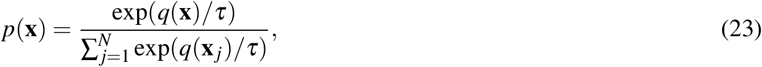

where *q*(**x**) is the predicted value of each option x for a given model (e.g., *q*(**x**) = UCB(**x**) for the UCB model), and *τ* is the temperature parameter. Lower values of *τ* indicate more concentrated probability distributions, corresponding to more precise predictions. All models include *τ* as a free parameter. Additionally, Function Learning models estimate *λ* (length-scale), Option Learning models estimate 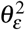 (error variance), and Upper Confidence Bound sampling models estimate *β* (exploration bonus).

#### Cross Validation.

We fit all models—per participant—using cross-validated MLE, with either a Differential Evolution algorithm^65^ or a grid search if the model contained only a single parameter. Parameter estimates are constrained to positive values in the range [exp(–5),exp(5)]. Cross-validation is performed by first separating participant data according to horizon length, which alternated between rounds (within subjects). For each participant, half of the rounds corresponded to a short horizon and the other half corresponded to a long horizon. Within all rounds of each horizon length, we use leave-one-out cross-validation to iteratively form a training set by leaving out a single round, computing a MLE on the training set, and then generating out-of-sample predictions on the remaining round. This is repeated for all combinations of training set and test set, and for both short and long horizon sets. The cross-validation procedure yielded one set of parameter estimates per round, per participant, and out-of-sample predictions for 120 choices in Experiment 1 and 240 choices in Experiments 2 and 3 (per participant).

#### Predictive Accuracy.

Prediction error (computed as log loss) is summed up over all rounds, and is reported as *predictive accuracy*, using a pseudo-*R*^2^ measure that compares the total log loss prediction error for each model to that of a random model:

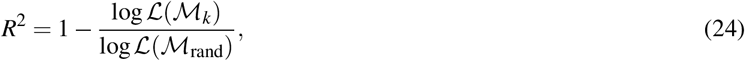

where log*𝓛*(*𝓜*_rand_) is the log loss of a random model (i.e., picking options with equal probability) and log*𝓛*(*𝓜*_*k*_) is the log loss of model *k*’s out-of-sample prediction error. Intuitively, *R*^2^ = 0 corresponds to prediction accuracy equivalent to chance, while *R*^2^ = 1 corresponds to theoretical perfect prediction accuracy, since log*𝓛*(*𝓜*_*k*_)/log*𝓛*(*𝓜*_rand_) → 0 when log*𝓛*(*𝓜*_*k*_) ≪ log*𝓛*(*𝓜*_rand_). *R*^2^ can also be below zero when the model predictions are worse than random chance.

### Simulated learning curves

We use participants’ cross-validated parameter estimates to specify a given model and then simulate performance. At each trial, model predictions correspond to a probabilistic distribution over options, which was then sampled and used to generate the observation for the next trial. In order to correspond with the manipulations of horizon length, payoff condition, and environment type, each simulation was performed at the participant level, producing data resembling a virtual participant for each replication. Iterating over each round, we selected the same environment as seen by the participant and then simulated data using the cross-validated parameters that were estimated using that round as the left-out round. Thus, just as model comparison was performed out-of-sample, the generated data was also out-of-sample, based on parameters that were estimated on a different set of rounds than the one being simulated. We performed 100 replications for each participant in each experiment, which were then aggregated to produce the learning curves in Figure 3b.

### Model Recovery

We present model recovery results that assess whether or not our predictive model comparison procedure allows us to correctly identify the true underlying model. To assess this, we generated data based on each individual participant’s parameter estimates (see above). We generated data using the Option Learning and the Function Learning Model for Experiment 1 and the Option Learning* Model and the Function Learning* Model for Experiments 2 and 3. In all cases, we used the UCB sampling strategy in conjunction with the specified learning model. We then utilized the same cross-validation method as before in order to determine if we could successfully identify which model generated the underlying data. Figure S2 shows the cross-validated predictive performance (half boxplot with each data point representing a single simulated participant) for the simulated data, along with the number of simulated participants best described (inset icon array).

**Figure S2.**
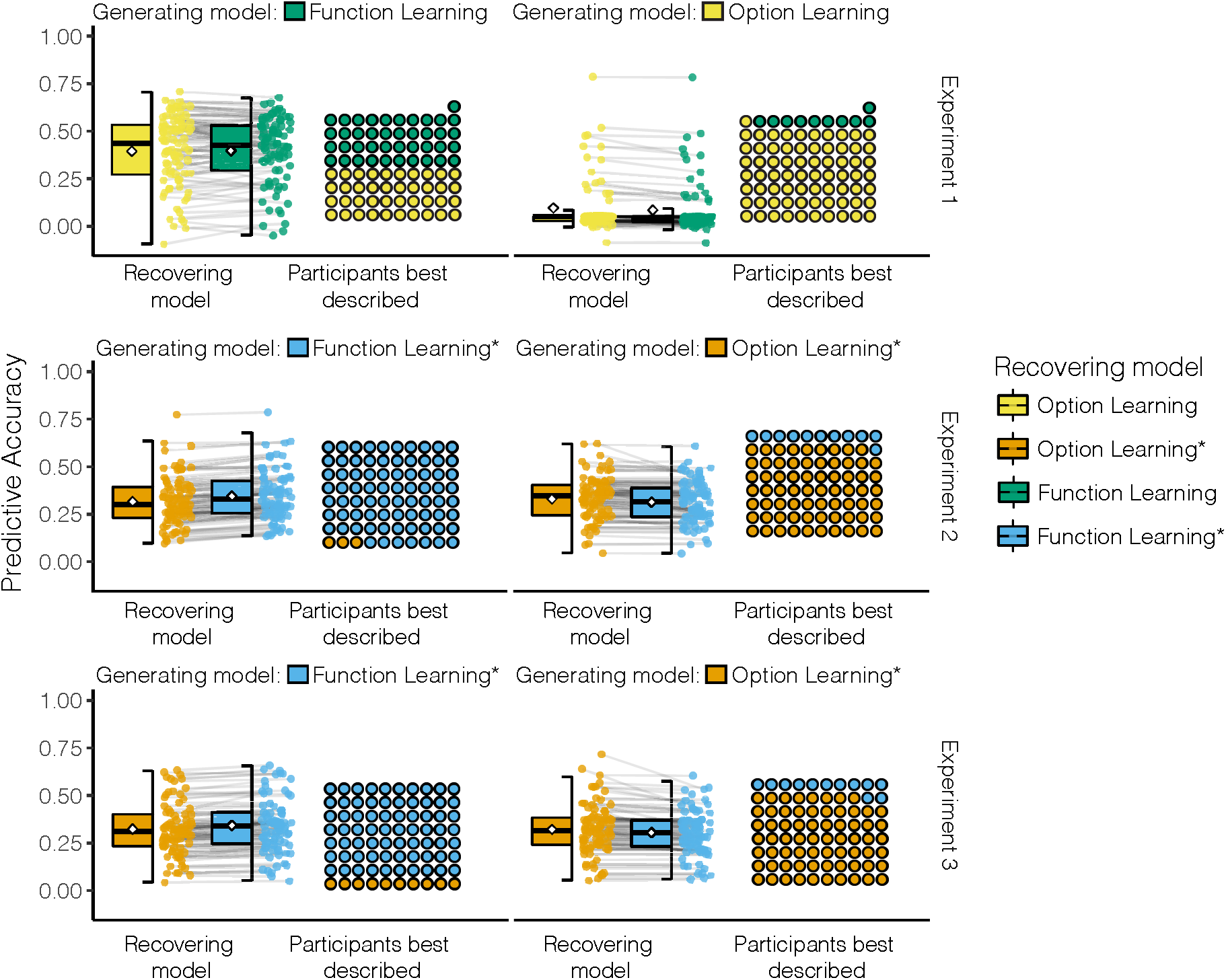
Model recovery results. Data was generated by the specified generating model (left and right columns) using individual participant parameter estimates. The recovery process used the same cross-validation method used in the model comparison. We report the predictive accuracy of each candidate recovery model (colours). Boxplots show the median (line), mean (diamond), interquartile range (box), and 1.5x IQR (whiskers). Each individual (simulated) participant is represented as a dot, with lines connecting each simulated participant. Icon arrays show the number of simulated participants best described. For both generating and recovery models, we used UCB sampling. Table S3 reports the median values of the cross-validated parameter estimates used to specify each generating model.

#### Experiment 1

In the simulation for Experiment 1, our predictive model comparison procedure shows that the Option Learning Model is a better predictor for data generated from the same underlying model, whereas the Function Learning model is only marginally better at predicting data generated from the same underlying model. This suggests that our main model comparison results are robust to Type I errors, and provides evidence that the better predictive accuracy of the Function Learning model for participant data is unlikely due to overfitting.

When the Function Learning Model has generated the underlying data, the same Function Learning Model achieves a predictive accuracy of *R*^2^ = .4 and describes 41 out of 81 simulated participants best, whereas the Option Learning model achieves a predictive accuracy of *R*^2^ = .39 and describes 40 participants best. Furthermore, the protected probability of exceedance for the Function Learning Model is pxp = 0.51. This makes our finding of the Function Learning Model as the best predictive model even stronger as, technically, the Option Learning Model could mimic parts of the Function Learning behaviour.

When the Option Learning Model generates data using participant parameter estimates, the same Option Learning Model achieves an average predictive accuracy of *R*^2^ = .1 and describes 71 out of 81 simulated participants best. On the same generated data, the Function Learning Model achieves an average predictive accuracy of *R*^2^ = .08 and only describes 10 out of 81 simulated participants best. The protected probability of exceedance for the Option Learning Model is pxp = 0.99. If the counterfactual had occurred, namely that if data generated by the Option Learning Model had been best predicted by the Function Learning Model, we would need to be sceptical about our modelling results on the basis that the wrong model could describe data better than the true generating model. However, here we see that the Function Learning Model does not make better predictions than the true model for data generated by the Option Learning Model.

#### Experiment 2

In the simulations for Experiment 2, we used the localized version of each type of learning model for both generation and recovery, since in both cases, localization improved model accuracy in predicting the human participants (Table S3). Here, we find very clear recoverability in all cases, with the recovering model best predicting the vast majority of simulated participants when it is also the generating model (Fig. S2).

When the Function Learning* Model generates the underlying data, the same Function Learning* Model achieves a predictive accuracy of *R*^2^ = .34 and describes 77 out of 80 simulated participants best, whereas the Option Learning* Model describes only 3 out of 80 simulated participants best, with a average predictive accuracy of *R*^2^ = .32. The protected probability of exceedance for the Function Learning* model is pxp = 1.

When the Option Learning* Model generates the data, the same Option Learning* Model achieves a predictive accuracy of *R*^2^ = .33 and predicts 69 out of 80 simulated participants best, whereas the Function Learning* Model predicts only 11 simulated participants best, with an average predictive accuracy of *R*^2^ = .31. The protected probability of exceedance for the Option Learning* model is pxp = 1. Again, we find evidence that the models are indeed discriminable, and that the Function Learning* Model does not overfit data generated by the wrong model.

#### Experiment 3

We again find in all cases the best recovery model is the same as the generating model. When the Function Learning* Model generates data, the matched recovery with the same Function Learning* Model best predicts 70 out of 80 participants, with an average predictive accuracy of *R*^2^ = .34. The Option Learning* Model best predicts the remaining 10 participants, with an average predictive accuracy of *R*^2^ = .32. The protected probability of exceedance for the Function Learning* model is pxp = 1.

When the Option Learning* Model generates the data, the same Option Learning* Model best predicts 68 out of 80 participants with an average predictive accuracy of *R*^2^ = .32, whereas the Function Learning* Model only best predicts 12 out of 80 participants with an average predictive accuracy of *R*^2^ = .3. The protected probability of exceedance for the Option Learning* model is pxp = 1.

In all simulations, the model that generates the underlying data is also the best performing model, as assessed by predictive accuracy, the number of simulated participants predicted best, and the protected probability of exceedance. Thus, we can confidently say that our cross-validation procedure distinguishes between these model classes. Moreover, in the cases where the Function Learning or Function Learning* Model generated the underlying data, the predictive accuracy of the same model is not perfect (i.e., *R*^2^ = 1), but rather close to the predictive accuracies we found for participant data (Table S3).

### High temperature recovery

We also assessed how much each model’s recovery can be affected by the underlying randomness of the softmax choice function. For every recovery simulation, we selected the 10 simulations with the highest underlying softmax temperature parameter *τ* (ranges: 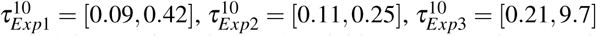) and again calculated the probability of exceedance for the true underlying model. The results of this analysis led to a probability of exceedance for the Function Learning Model in Experiment 1 of pxp = .81, for the Function Learning* Model in Experiment 2 of pxp = 0.99, for the Function Learning* Model in Experiment 3 of pxp = 0.93, for the Option Learning Model in Experiment 1 of pxp = 0.97, for the Option Learning* Model in Experiment 2 of pxp = 0.99, and for the Option Learning Model in Experiment 3 of pxp = 0.98. Thus, the models seem to be well-recoverable even in scenarios with high levels of random noise in the generated responses.

### Parameter Recovery

Another important question is whether or not the reported parameter estimates of the two Function Learning models are reliable and robust. We address this question by assessing the recoverability of the three parameters of the Function Learning model, the length-scale *λ*, the exploration factor *β*, and the temperature parameter *τ* of the softmax choice rule. We use the results from the model recovery simulation described above, and correlate the empirically estimated parameters used to generate data (i.e., the estimates based on participants’ data), with the parameter estimates of the recovering model (i.e., the MLE from the cross-validation procedure on the simulated data). We assess whether the recovered parameter estimates are similar to the parameters that were used to generated the underlying data. We present parameter recovery results for the Function Learning Model for Experiment 1 and the Function Learning* Model for Experiments 2 and 3, in all cases using the UCB sampling strategy. We report the results in Figure S3, with the generating parameter estimate on the x-axis and the recovered parameter estimate on the y-axis. We report rank-correlation using Kendall’s tau (*r*_*τ*_), which should not be confused with the temperature parameter *τ* of the softmax function. Additionally, we calculate the Bayes Factor (*BF*_*τ*_) to quantify the evidence for the presence of a positive correlation using non-informative, shifted, and scaled beta-priors as recommended by^66^.

**Figure S3.**
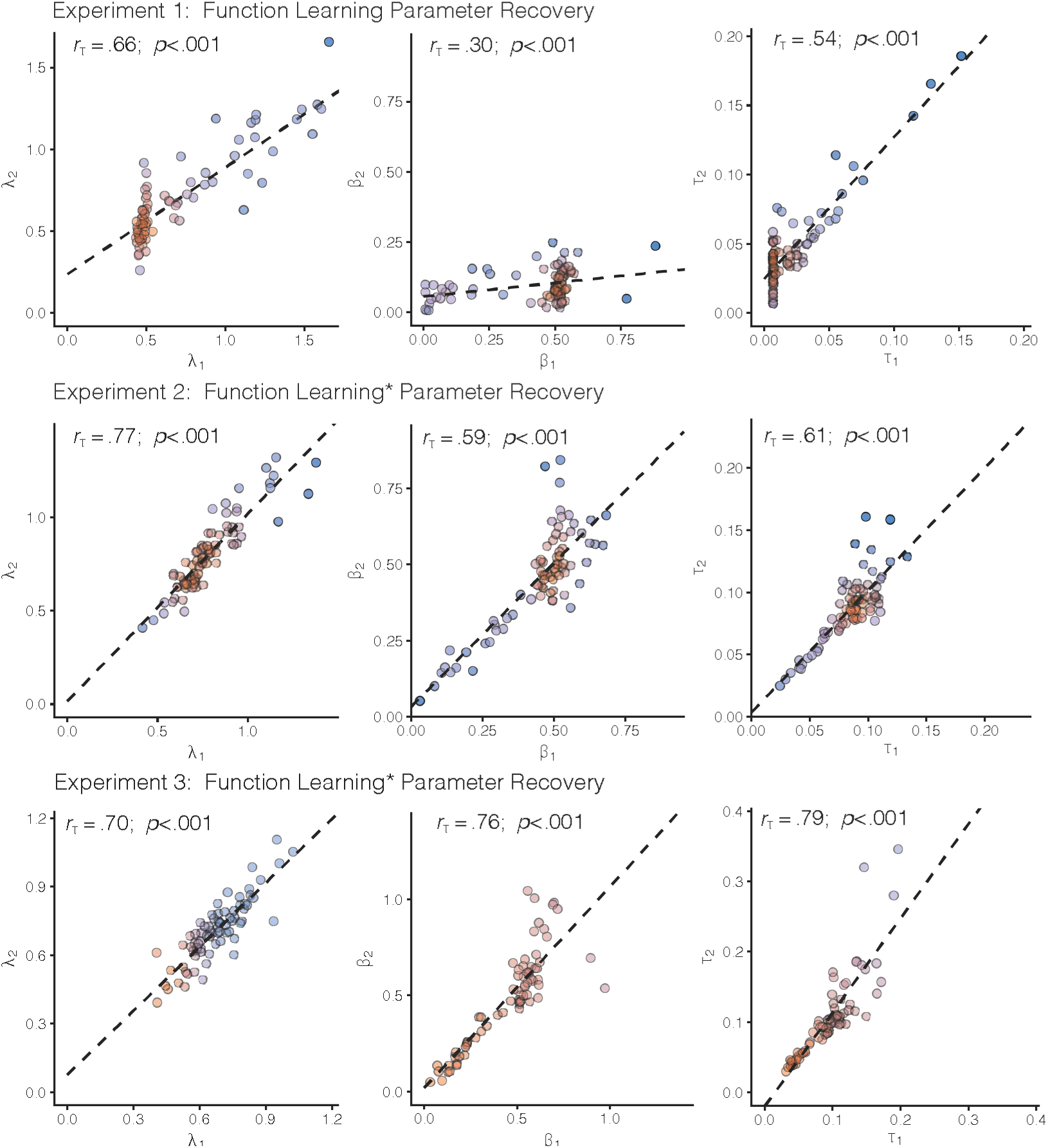
Parameter recovery. The generating parameter estimate is on the x-axis and the recovered parameter estimate is on the y-axis. The generating parameter estimates are from the cross-validated participant parameter estimates, which were used to simulate data. Recovered parameter estimates are the result of the cross-validated model comparison on the simulated data. While the cross-validation procedure yielded *k* estimates per participant, one for each round (*k*_*Exp*1_ = 16; *k*_*Exp*2_ = *k*_*Exp*3_ = 8), we show the median estimate per (simulated) participant. The dashed line shows a linear regression on the data, with the rank correlation (Kendall’s tau) and p-value shown above. For readability, colours represent the bivariate kernel density estimate, with red indicating higher density. The axis limits are chosen based on 1.5× the IQR for the larger of the two values (generating or recovered parameter estimates). Thus, some outliers are omitted from these plots (2.3% in Exp. 1, 1.7% in Exp. 2, and 5.2% in Exp. 3) but all datapoints are used to calculate the rank correlations.

For Experiment 1, the rank-correlation between the generating and the recovered length-scale *λ* is *r*_*τ*_ = .66, *p* < .001, *BF*_*τ*_ > 100, the correlation between the generating and the recovered exploration factor *β* is *r*_*τ*_ = .30, *p* < .001, *BF*_*τ*_ > 100, and the correlation between the generating and the recovered softmax temperature parameter *τ* is *r*_*τ*_ = .54, *p* < .001, *BF*_*τ*_ > 100. For Experiment 2, the correlation between the generating and the recovered *λ* is *r*_*τ*_ = .77, *p* < .001, *BF*_*τ*_ > 100, for *β* the correlation is *r*_*τ*_ = .59, *p* < .001, *BF*_*τ*_ > 100, and for *τ* the correlation is *r* =_*τ*_ .61, *p* < .001, *BF*_*τ*_ > 100. For Experiment 3, the correlation between the generating and the recovered *λ* is *r*_*τ*_ = .70, *p* < .001, *BF*_*τ*_ > 100, for *β* the correlation is *r*_*τ*_ = .76, *p* < .001, *BF*_*τ*_ > 100, and for *τ* the correlation is *r* = .79, *p* < .001, *BF*_*τ*_ > 100.

These results show that the rank-correlation between the generating and the recovered parameters is very high for all experiments and for all parameters. Thus, we have strong evidence to support the claim that the reported parameter estimates of the Function Learning Model (Table S3) are reliable, and therefore interpretable. Importantly, we find that estimates for *β* (exploration bonus) and *τ* (softmax temperature) are indeed separately identifiable, providing evidence for the existence of a *directed* exploration bonus^12^, as a separate phenomena from noisy, undirected exploration^47^ in our data.

### Experimental conditions and model characteristics

To further assess how the experimental conditions influenced the model’s behaviour, we performed Bayesian linear regressions of the experimental conditions onto the models’ predictive accuracy and parameter estimates. To do so, we assumed a Gaussian prior on the coefficients, and an inverse Gamma prior on the conditional error variance, while inference was performed via Gibbs sampling. The results of these regressions are shown in Table S1. Whereas the smoothness of the underlying environments (in Experiments 1 and 2) had no effect on the model’s predictive accuracy and almost no effect on parameter estimates (apart from a small effect on directed exploration in Experiment 1), participants in the Accumulation payoff condition showed decreased levels of directed exploration (as captured by *β*) in Experiment 1 and Experiment 3, and decreased levels of random exploration in Experiment 3. Thus, our model seems to capture meaningful differences between the two reward conditions in these two experiments.

### Mismatched generalization

#### Generalized mismatch

A mismatch is defined as estimating a different level of spatial correlations (captured by the per participant *λ*-estimates) than the ground truth in the environment. In the main text (Fig. 4), we report a generalized Bayesian optimization simulation where we simulate every possible combination between *λ*_0_ = {0.1,0.2,···,1} and *λ*_1_ = {0.1,0.2,···,1}, leading to 100 different combinations of student-teacher scenarios. For each of these combinations, we sample a continuous bivariate target function from a GP parameterized by *λ*_0_ and then use the Function Learning-UCB Model parameterized by *λ*_1_ to search for rewards. The exploration parameter *β* was set to 0.5 to resemble participant behaviour (Table S3). The input space was continuous between 0 and 1, i.e., any number between 0 and 1 could be chosen and GP-UCB was optimized (sometimes called the inner-optimization loop) per step using NLOPT^81^ for non-linear optimization. It should be noted that instead of using a softmax choice rule, the optimization method uses an argmax rule, since the former is not defined for continuous input spaces. Additionally, since the interpretation of *λ* is always relative to the input range, a length-scale of *λ* = 1 along the unit input range would be equivalent to *λ* = 10 in the *x*,*y* = [0,10] input range of Experiments 2 and 3. Thus, this simulation represents a broad set of potential mismatch alignments, while the use of continuous inputs extends the scope of the task to an infinite state space.

##### Experiments 1 and 2

In both Experiments 1 and 2, we found that participant *λ*-estimates were systematically lower than the true value (*λ*_*Rough*_ = 1 and *λ*_*Smooth*_ = 2), which can be interpreted as a tendency to undergeneralize compared to the spatial correlation between rewards. In order to test how this tendency to undergeneralize (i.e., underestimate *λ*) influences task performance, we conducted two additional sets of simulations using the exact experimental design for Experiments 1 and 2 (Fig. S4a-b). These simulations used different combinations of *λ* values in a *teacher* kernel (x-axis) to generate environments and in a *student* kernel (y-axis), to simulate human search behaviour with the Function Learning Model.

Both teacher and student kernels were always RBF kernels, where the teacher kernel (used to generate environments) was parameterized with a length-scale *λ*_0_ and the student kernel (used to simulate search behaviour) with a length-scale *λ*_1_. For situations in which *λ*_0_ ≠ *λ*_1_, the assumptions of the student can be seen as mismatched with the environment. The student *overgeneralizes* when *λ*_1_ > *λ*_0_ (Fig. S4a-b above the dotted line), and *undergeneralizes* when *λ*_1_ > *λ*_0_ (Fig. S4a-b below the dotted line), as was captured by our behavioural data. We simulated each possible combination of *λ*_0_ = {0.1,0.2,···,3} and *λ*_1_ = {0.1,0.2,···,3}, leading to 900 different combinations of student-teacher scenarios. For each of these combinations, we sampled a target function from a GP parameterized by *λ*_0_ and then used the Function Learning-UCB Model parameterized by *λ*_1_ to search for rewards using the median parameter estimates for *β* and *τ* from the matching experiment (see Table S3).

Figures S4a-b show the results of the Experiment 1 and Experiment 2 simulations, where the colour of each tile shows the median reward obtained at the indicated trial number, for each of the 100 replications using the specified teacher-student scenario. The first simulation assessed mismatch in the univariate setting of Experiment 1 (Fig. S4a), using the median participant estimates of both the softmax temperature parameter *τ* = 0.01 and the exploration parameter *β* = 0.50 and simulating 100 replications for every combination between *λ*_0_ = {0.1,0.2,···,3} and *λ*_1_ = {0.1,0.2,···,3}. This simulation showed that it can be beneficial to undergeneralize (Fig. S4a, area below the dotted line), in particular during the first five trials. Repeating the same simulations for the bivariate setting of Experiment 2 (using the median participant estimates *τ* = 0.02 and *β* = 0.47), we found that undergeneralization can also be beneficial in a more complex two-dimensional environment (Fig. S4b), at least in the early phases of learning. In general, assumptions about the level of correlations in the environment (i.e., extent of generalization *λ*) only influence rewards in the short term, and can disappear over time once each option has been sufficiently sampled^25^.

**Figure S4.**
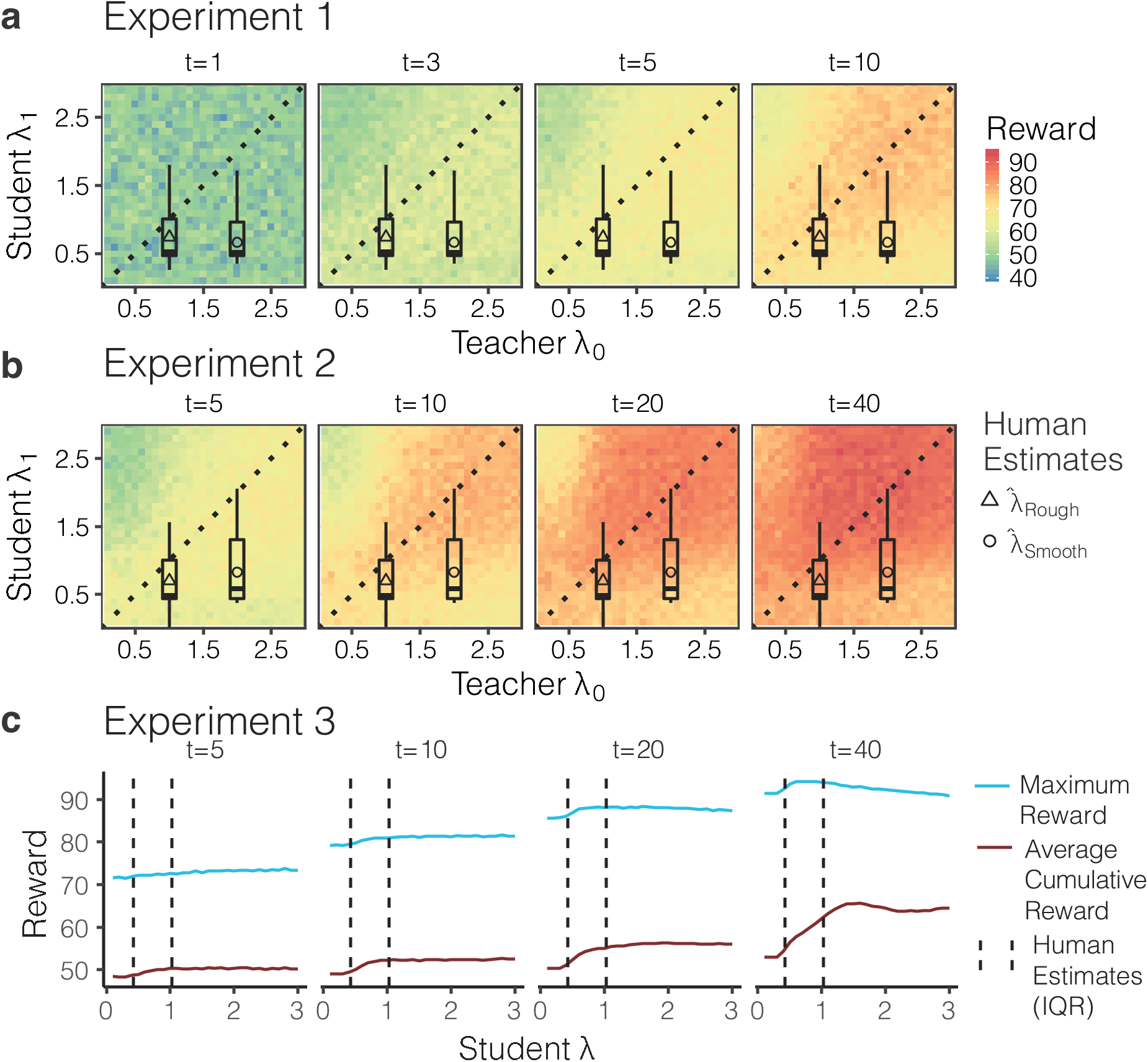
Mismatched length-scale (*λ*) simulation results. **a-b)** The teacher length-scale *λ*_0_ is on the x-axis, the student length-scale *λ*_1_ is on the y-axis, and each panel represents a different trial *t*. The teacher *λ*_0_ values were used to generate environments, while the student *λ*_1_ values were used to parameterize the Function Learning-UCB Model to simulate search performance. The dotted lines show where *λ*_0_ = *λ*_1_ and mark the difference between undergeneralization and overgeneralization, with points below the line indicating undergeneralization. Each tile of the heat-map indicates the median reward obtained for that particular *λ*_0_-*λ*_1_-combination, aggregated over 100 replications. Triangles and circles indicate mean participant *λ* estimates from Rough and Smooth conditions, with boxplots showing the interquartile range, the median (line), and 1.5x IQR (whiskers). c) Simulations with student *λ* values in the range [0,3] over 10,000 samples (sampled with replacement) from the set of 20 different natural environments. Red lines show average cumulative reward and blue lines show the maximum reward. Vertical dashed lines show the interquartile range of participant *λ* estimates.

##### Experiment 3

Given the robust tendency to undergeneralize in Experiments 1 and 2 (where there was a true underlying level of spatial correlation), we ran one last simulation to examine how adaptive participant *λ* estimates were in the real-world datasets used in Experiment 3, compared to other possible *λ* values. Figure S4c shows the performance of different student *λ* values in the range {0.1,0.2,···,3} simulated over 10,000 replications sampled (with replacement) from the set of 20 natural environments. Red lines show performance in terms of average cumulative reward (Accumulation criterion) and blue lines show performance in terms of maximum reward (Maximization criterion). Vertical dashed lines indicate the interquartile range of participant *λ* estimates. As student *λ* values increase, performance by both metrics typically peaks within the range of human *λ* estimates, with performance largely staying constant or decreasing for larger levels of *λ* (with the exception of average reward at *τ* = 40). Thus, we find that the extent of generalization observed in participants is generally adaptive to the real-world environments they encountered. It should also be noted that higher levels of generalization beyond what we observed in participant data have only marginal benefits, yet could potentially come with additional computational costs (depending on how it is implemented). Recall that a *λ* of 1 corresponds to assuming the correlation of rewards effectively decays to 0 for options with a distance greater than 3. If we assume a computational implementation where information about uncorrelated options is disregarded (e.g., in a sparse GP^67^), then the range of participant *λ* estimates could suggest a tendency towards lower complexity and memory requirements, while sacrificing only marginal benefits in terms of either average cumulative reward or maximum reward.

### Natural Environments

The environments used in Experiment 3 were compiled from various agricultural datasets^34, 68–80^ (Table S2), where payoffs correspond to normalized crop yield (by weight), and the rows and columns of the 11x11 grid correspond to the rows and columns of a field. Because agricultural data is naturally discretized into a grid, we did not need to interpolate or transform the data in any way (so as not to introduce any additional assumptions), except for the normalization of payoffs in the range [0,100], where 0 corresponds to the lowest yield and 100 corresponds to the largest yield. Note that as in the other experiments, Gaussian noise was added to each observed payoff in the experiment.

In selecting datasets, we used three inclusion criteria. Firstly, the datasets needed to be at least as large as our 11x11 grid. If the dataset was larger, we randomly sampled a 11x11 subsection from the data. Secondly, to avoid datasets where payoffs were highly skewed (e.g., with the majority of payoffs around 0 or around 100), we only included datasets where the median payoff was in the range [25,75]. Lastly, we required that the spatial autocorrelation of each environment (computed using Moran’s *I*) be positive:

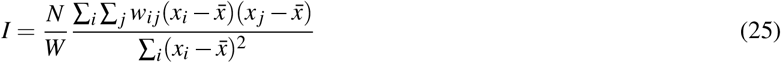

where *N* is the total number of samples (i.e., each of the 121 sections of land in a 11×11 grid), *x*_*i*_ is the normalized yield (i.e., payoff) for option *i*,*x̄* is the mean payoff over all samples, and *W* is the spatial weights matrix where *w*_*ij*_ = 1 ~ *i* and *j* are the same or neighbouring samples and *w*_*ij*_ = 0 otherwise. Moran’s *I* ranges between [–1,1] where intuitively *I* = –1 would resemble a checkerboard pattern (with black and white tiles reflecting the highest and lowest values in the payoff spectrum), indicating maximum difference between neighbouring samples. On the other hand, *I* → 1 would reflect a linear step function, with maximally high payoffs on one side of the environment and maximally low payoffs on the other side. We included all environments where *I* > 0, indicating that there exists some level of positive spatial correlation that could be used by participants to guide search.

Although the structure of rewards in real-world data can sometimes be distributed differently and in particular more discretely (for example, imagine a bitmap or other structural patterns such as a checkerboard or a crop circle), we believe that our environment inclusion criteria allow us to appropriately model generalization using our pool of models, while at the same time extending the scope to more complex and challenging natural structures.

### Additional Behavioural Analyses

#### Learning over trials and rounds

We assessed whether participants improved more strongly over trials or over rounds (Fig. S5). If they improved more over trials, this means that they are indeed finding better and better options, whereas if they are improving over rounds, this would also suggest some kind of meta-learning as they would get better at the task the more rounds they have performed previously. To test this, we fit a linear regression to every participant’s outcome individually, either only with trials or only with rounds as the independent variable. Afterwards, we extract the mean standardized slopes for each participant including their standard errors. Notice that these estimates are based on a linear regression, whereas learning curves are probably non-linear. Thus, this method might underestimate the true underlying effect of learning over time.

**Figure S5.**
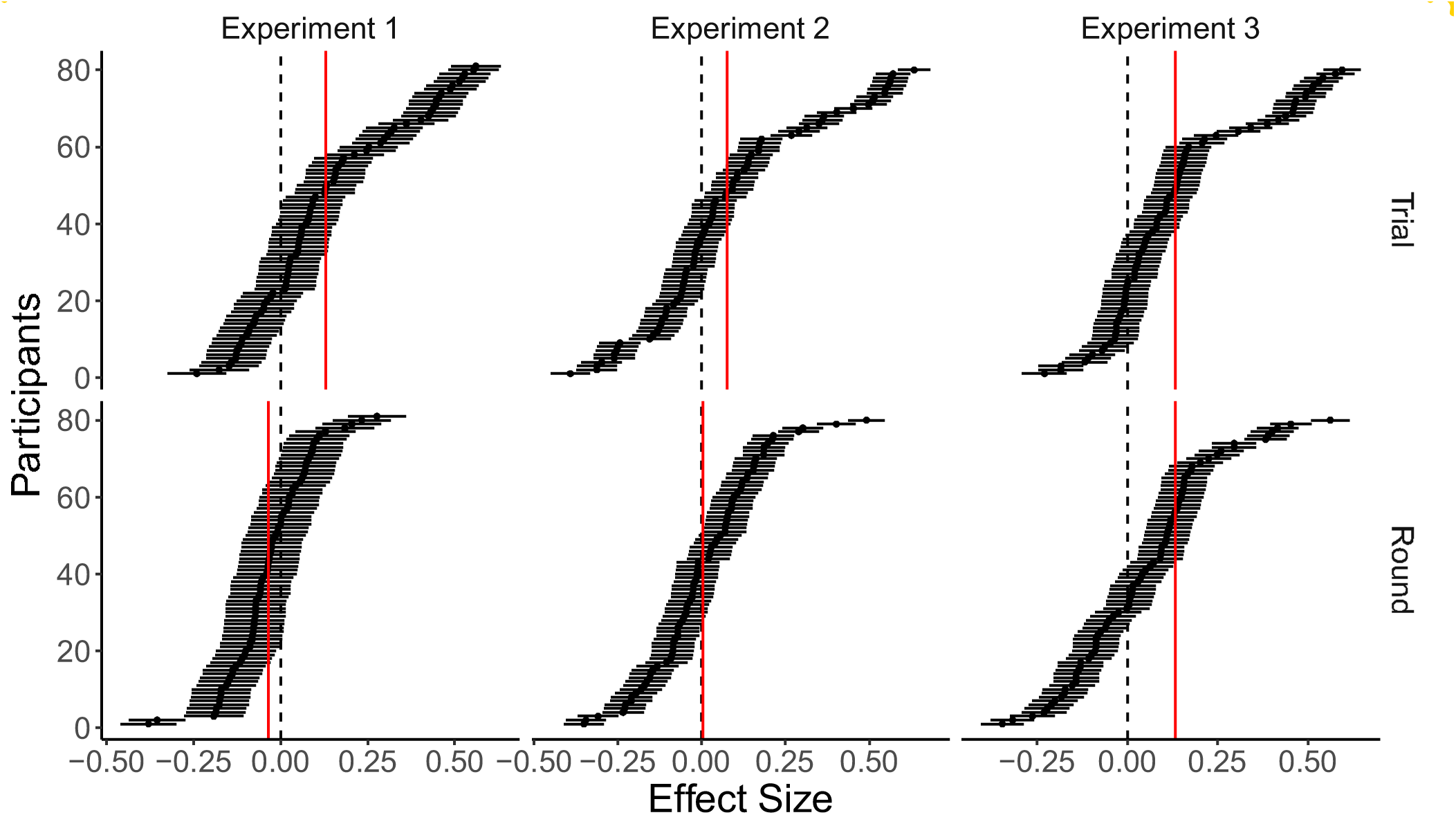
Learning over trials and rounds. Average correlational effect size of trial and round on score per participant as assessed by a standardized linear regression. Participants are ordered by effect size in decreasing order. Dashed lines indicate no effect. Red lines indicate average effect size.

Results (from one-sample *t*-tests with *μ*_0_ = 0) show that participants’ scores improve significantly over trials for Experiment 1 (*t*(80) = 5.57, *p* < .001, *d* = 0.6, 95% CI (0.2,1.1), *BF* > 100), Experiment 2 (*t*(79) = 2.78, *p* < .001, *d* = 0.31, 95% CI (–0.1,0.8), *BF* = 4.4), and Experiment 3 (*t*(79) = 5.91, *p* < .001, *d* = 0.7, 95% CI (0.2,1.1), *BF* > 100). Over successive rounds, there was a negative influence on performance in Experiment 1 (*t*(80) = –2.78, *p* = .007, *d* = –0.3, 95% CI (–0.7,0.1), *BF* = 4.3), no difference in Experiment 2 (*t*(79) = 0.21, *p* = .834, *d* = 0.02, 95% CI (–0.4,0.5), *BF* = 0.1), and a minor positive influence in Experiment 3 (*t*(79) = 2.16, *p* = .034, *d* = 0.2, 95% CI (–0.2,0.7), *BF* = 1.1). Overall, participants robustly improved over trials in all experiments, with the largest effect sizes found in Experiments 1 and 3. There was no improvement over rounds in all of the experiments, suggesting that the four fully revealed example environments presented prior to the start of the task was sufficient for familiarizing participants with the task.

#### Individual Learning Curves

To better understand why the aggregated participant learning curves sometimes decrease in average reward over time, whereas the simulated model curves tend not to (Fig. 3b), we present individual participant learning curves in Figure S6. Here, we separate the behavioural data by horizon (colour), payoff condition (rows), and environment (columns), where each line represents a single participant. We report performance in terms of both average reward (top section: Accumulation goal) and maximum reward (bottom section: Maximization goal).

**Figure S6.**
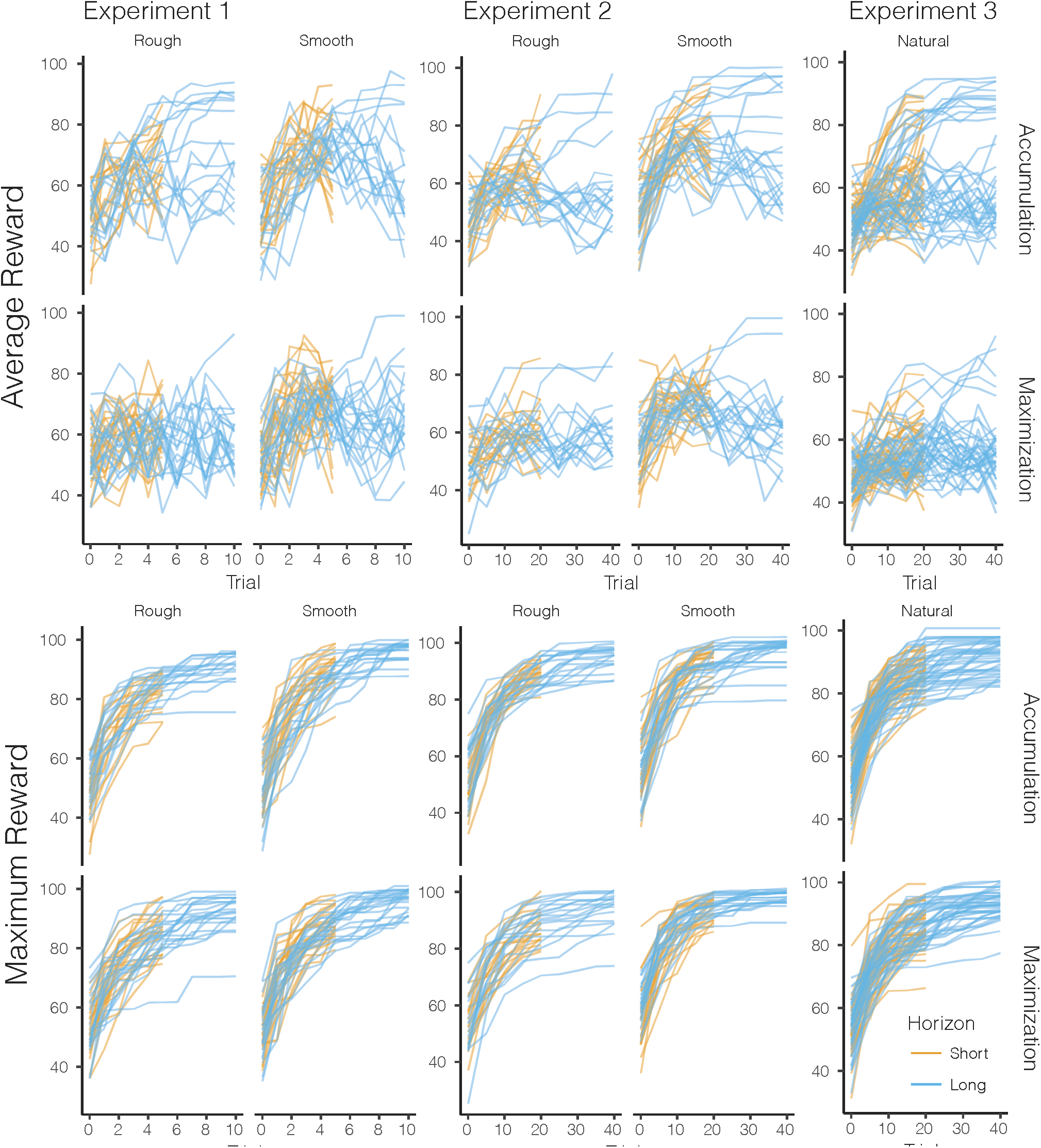
Individual participant learning curves. Each line represents a single participant, separated by search horizon (colour), by payoff condition (rows), and environment (columns). The top section shows performance in terms of average reward, while the bottom section shows performance in terms of maximum reward.

The individual learning curves reveal two main causes for the decrease in reward over time when aggregating over conditions and participants. Firstly, looking at the learning curves for participants assigned to the Accumulation condition (Fig. S6 top row), we see that roughly half of participants in the long search horizon (blue lines) show a decreasing trend at the midway point of the round. However, the other half of participants continue to gain increasingly higher rewards, more like the simulated learning curves of the Function Learning model in Figure 3b. This may be a by-product of the alternating search horizon manipulation, since the curves typically tend to decrease near the trial where a short horizon round would have ended, but also a tendency towards over-exploration that more closely resembles the Maximization goal.

Secondly, in aggregating over conditions and participants, the performance of the Accumulation and Maximization participants are averaged together. Whereas many Accumulation payoff condition participants display more positively increasing average reward, these data points are washed out by the Maximization payoff condition participants who tend to have flatter average reward curves in pursuit of the global optimization goal.

Lastly, one additional insight from the individual learning curves comes from the flat-lined maximum reward lines (S6, bottom section). Found more often in Accumulation participants, these flat lines represent participants who have reached a satisfactory payoff and cease additional exploration in order to exploit it. This is yet another behavioural signature of the payoff manipulations.

### Experiment Instructions

Figures S7-S9 provide screenshots from each experiment, showing the instructions provided to participants, separated by payoff condition. The top row of each figure shows the initial instructions, while the bottom row shows a set of summarized instructions provided alongside the task. Links to each of the experiments are also provided below.

- **Experiment 1:** https://arc-vlab.mpib-berlin.mpg.de/wu/gridsearch1/experiment1.html
- **Experiment 2:** https://arc-vlab.mpib-berlin.mpg.de/wu/gridsearch2/experiment2.html
- **Experiment 3:** href="https://arc-vlab.mpib-berlin.mpg.de/wu/gridsearch3/experiment3.html

**Figure S7.**
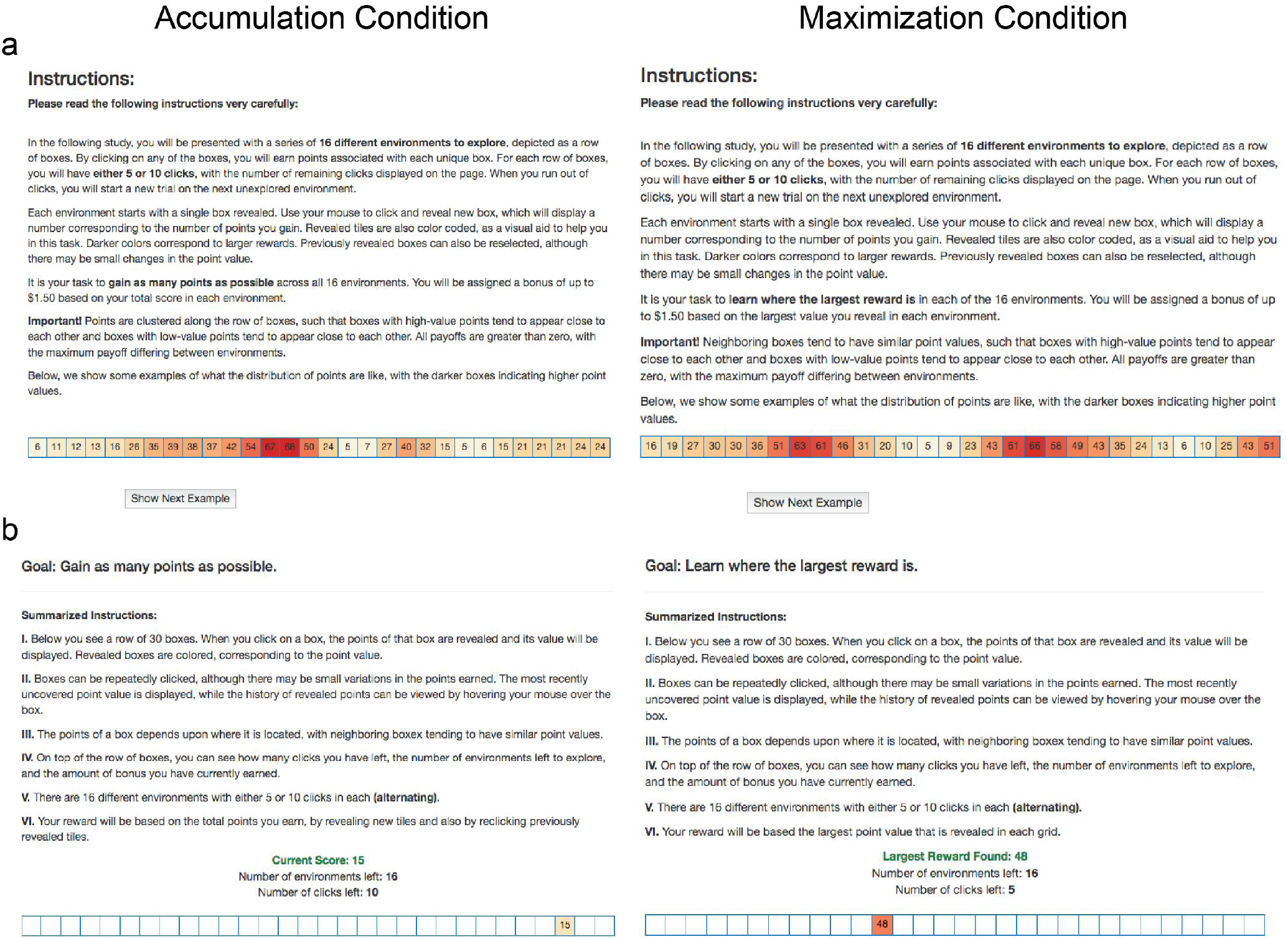
Screenshots from Experiment 1. Accumulation condition on the left and Maximization condition on the right. **a)** Initial instructions given to participants, followed by **b**) summarized instructions provided alongside the task.

**Figure S8.**
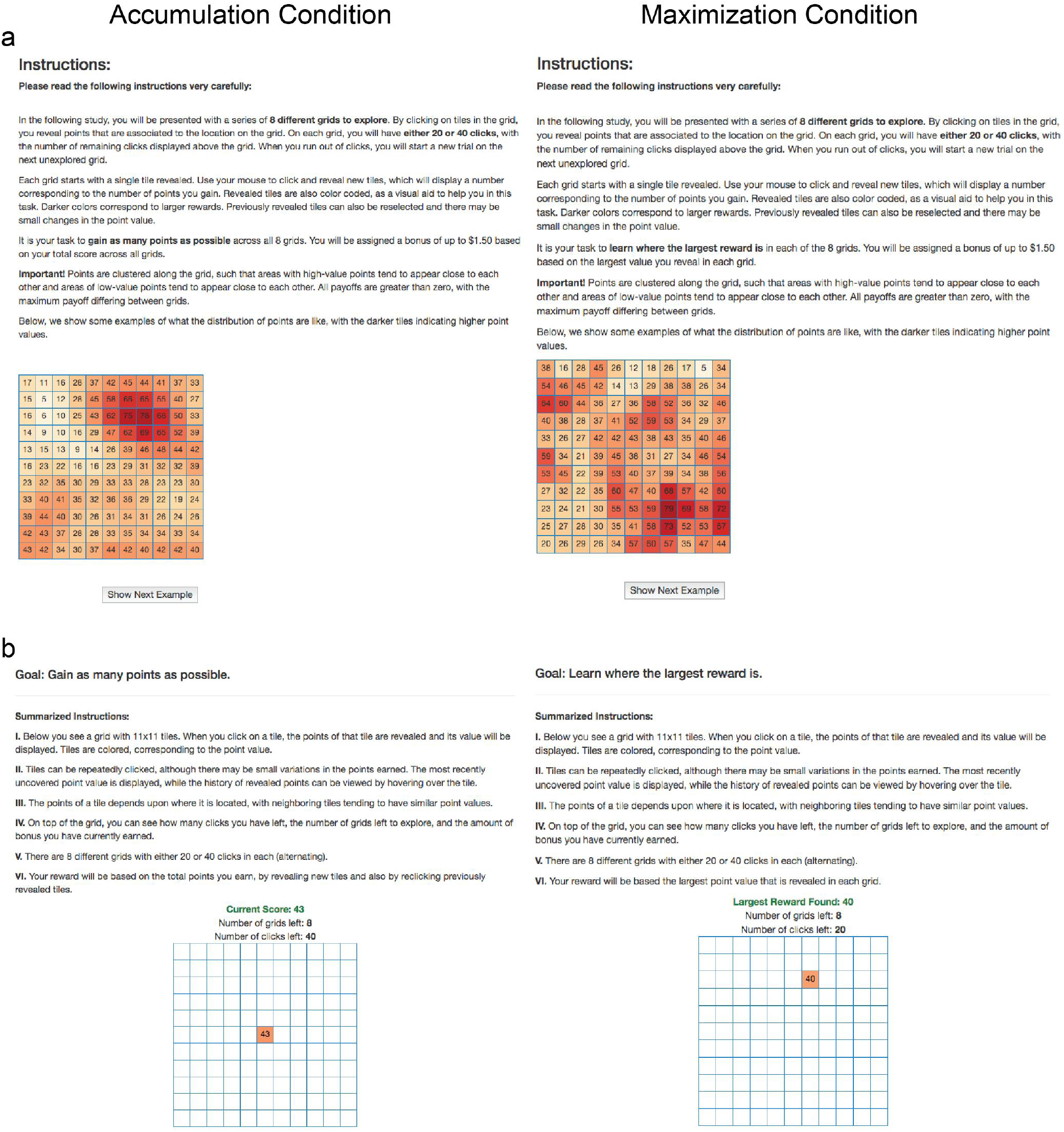
Screenshots from Experiment 2. Accumulation condition on the left and Maximization condition on the right. **a**) Initial instructions given to participants, followed by **b**) summarized instructions provided alongside the task.

**Figure S9.**
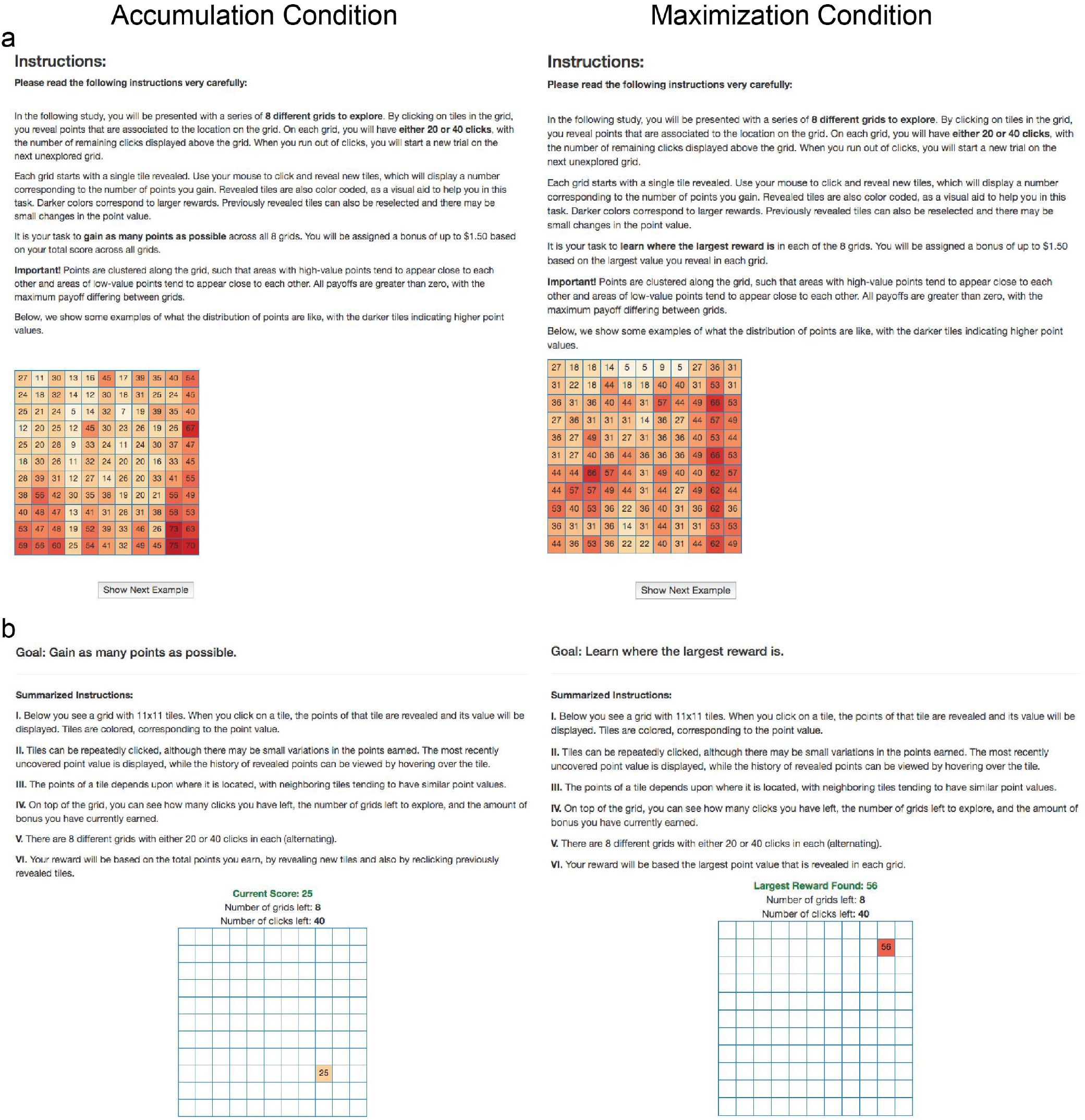
Screenshots from Experiment 3. Accumulation condition on the left and Maximization condition on the right. **a**) Initial instructions given to participants, followed by **b**) summarized instructions provided alongside the task.

**Table 1.**
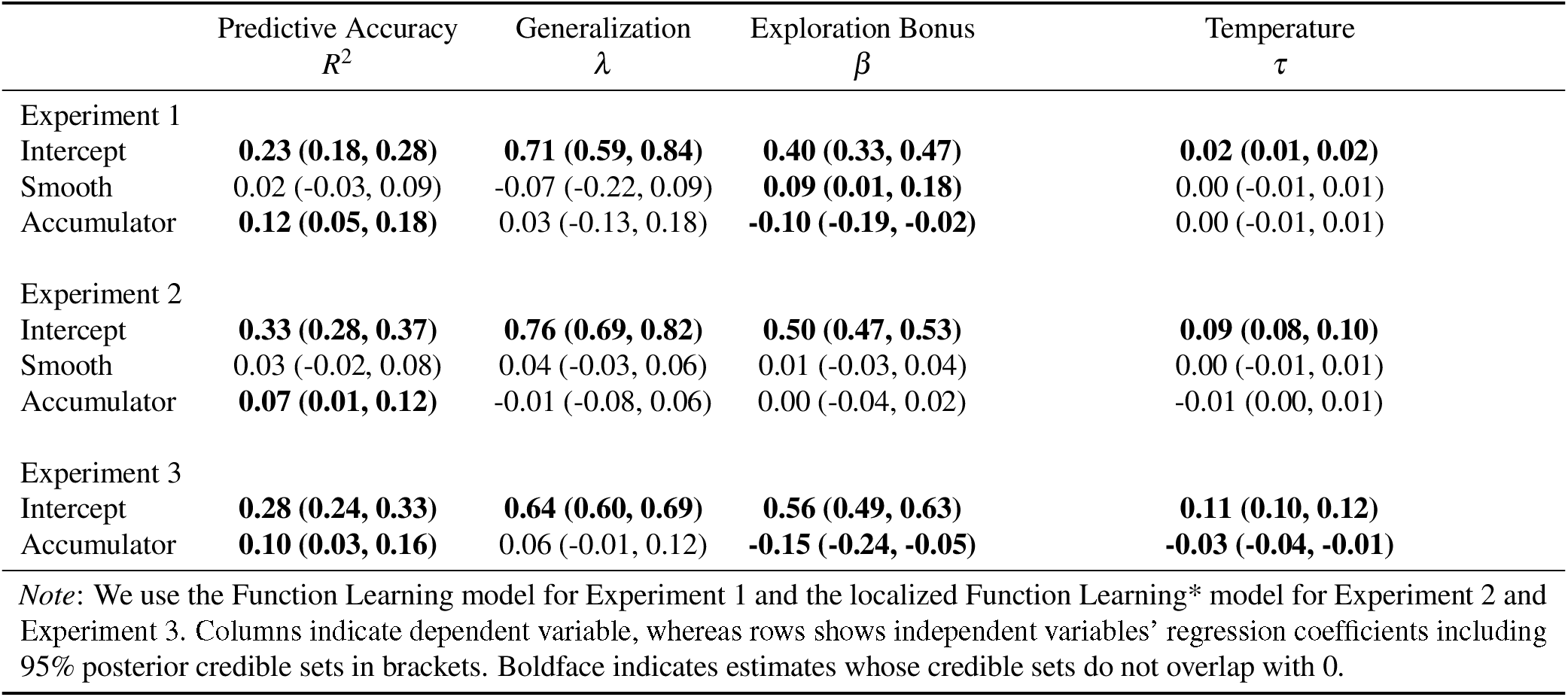
Bayesian linear regression of experimental conditions on model performance and parameter estimates.

**Table 2.**
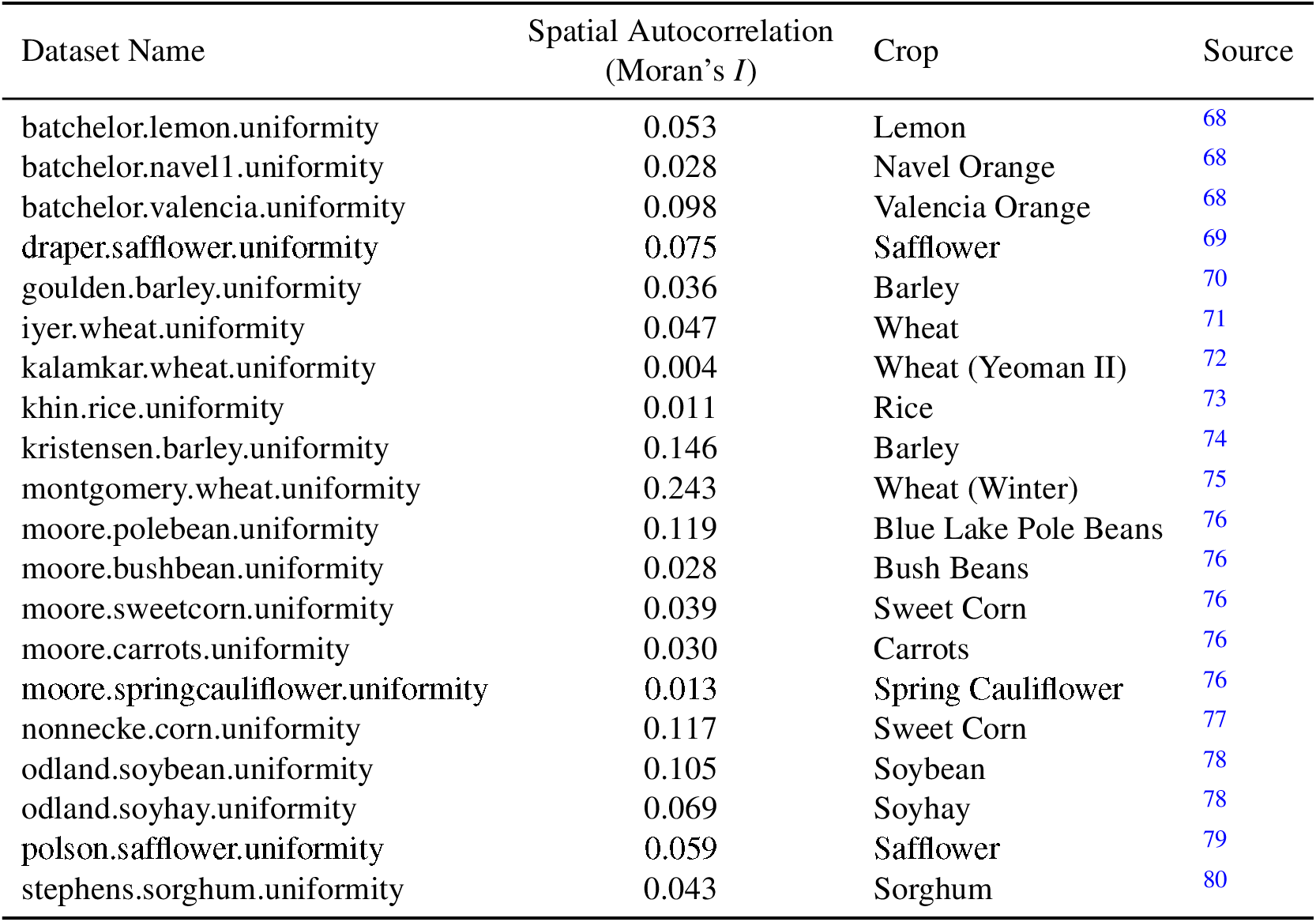
Agricultural datasets used in Experiment 3

**Table 3.**
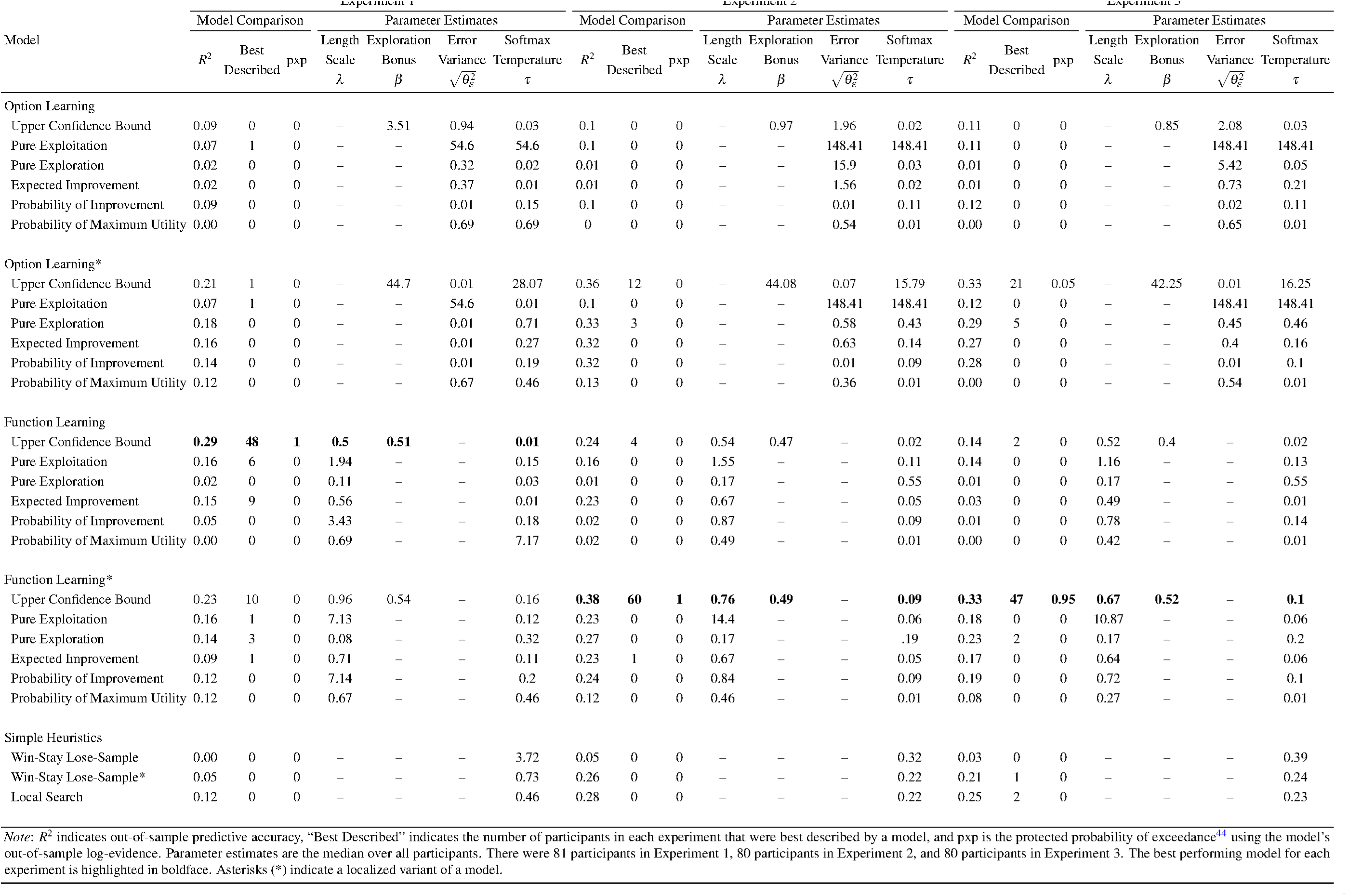
Modelling Result

